# Covalent Modification and Regulation of the Nuclear Receptor Nurr1 by a Dopamine Metabolite

**DOI:** 10.1101/368399

**Authors:** John M. Bruning, Yan Wang, Francesca Oltrabella, Boxue Tian, Svetlana A. Kholodar, Harrison Liu, Paulomi Bhattacharya, Su Guo, James M. Holton, Robert J. Fletterick, Matthew P. Jacobson, Pamela M. England

## Abstract

Nurr1, a nuclear receptor essential for the development, maintenance, and survival of midbrain dopaminergic neurons, is a potential therapeutic target for Parkinson’s disease, a neurological disorder characterized by the degeneration of these same neurons. Efforts to identify Nurr1 agonists have been hampered by the recognition that it lacks several classic regulatory elements of nuclear receptor function, including the canonical ligand-binding pocket. Here we report that the dopamine metabolite 5,6-dihydroxyindole (DHI) binds directly to and modulates the activity of Nurr1. Using biophysical assays and x-ray crystallography we show that DHI binds to the ligand binding domain within a non-canonical pocket, forming a covalent adduct with Cys566. In cultured cells and zebrafish, DHI stimulates Nurr1 activity, including the transcription of target genes underlying dopamine homeostasis. These findings suggest avenues for developing synthetic Nurr1 ligands to ameliorate the symptoms and progression of Parkinson’s disease.

## INTRODUCTION

Parkinson’s disease (PD) is a neurological disorder afflicting 10 million people worldwide (Wirdefeldt et al., 2011). In an estimated 90% of PD patients, the cause of the disease is unknown, having no clear genetic or environmental origin (de Lau and Breteler, 2006). The most pronounced neuropathological feature of PD is the progressive degeneration of dopaminergic neurons in the substantia nigra pars compacta and the consequent reduction in dopamine levels in the striatum, which manifest as impairments in motor function (e.g. rigidity, tremor, bradykinesia) (Samii et al., 2004). Notably, this degeneration appears to be *preceded* by the loss of the dopaminergic phenotype; that is, at least some dopaminergic neurons first stop producing and signaling with dopamine prior to degenerating (Janezic et al., 2013). Although the molecular basis for idiopathic PD remains incompletely understood, it has been proposed to include oxidative stress, mitochondrial dysfunction, and dysregulation of dopamine homeostasis (Blesa et al., 2015; Hauser and Hastings, 2013; Hwang, 2013). Currently, there are no available treatments that stop or even slow the progression of PD. Existing therapeutics relieve PD symptoms by increasing dopaminergic signaling through one of three mechanisms: (1) increasing dopamine levels by augmenting the amount of its biosynthetic precursor, L-DOPA; (2) blocking the breakdown of dopamine by inhibiting its metabolic enzymes (MAO, COMT); (3) mimicking the activity of dopamine by directly agonizing dopamine receptors. However, these drugs only partially alleviate symptoms and can have significant side effects, especially as the disease progresses. New types of therapeutics are desperately needed to combat both the symptoms and progression of PD.

The nuclear receptor related-1 protein, Nurr1 (NR4A2), is a transcription factor that regulates the expression of genes critical for the development, maintenance, and survival of dopaminergic neurons (Alavian et al., 2014; Decressac et al., 2013; Dong et al., 2016; Jankovic et al., 2005; Johnson et al., 2011; Kadkhodaei et al., 2009; Luo, 2012; Zetterstrom et al., 1997). In particular, Nurr1 plays a fundamental role in maintaining dopamine homeostasis by regulating transcription of the genes governing dopamine synthesis (*TH*, tyrosine hydroxylase; *DDC*, dopa decarboxylase), packaging (*SLC18A2*, vesicular monoamine transporter 2, VMAT2), and reuptake (*DAT*, dopamine transporter, also known as *SLC6A3*)(Hermanson et al., 2003; Iwawaki et al., 2000; Johnson et al., 2011; Sacchetti et al., 2001) (Figure 1A). Nurr1 also regulates the survival of dopaminergic neurons by stimulating the transcription of genes coding for neurotrophic factors (BDNF, NGF), anti-inflammatory responses (GDNF receptor c-Ret), and oxidative stress management (SOD1), as well as repressing the transcription of pro-inflammatory genes (TNFalpha, iNOS, IL-1beta) (Galleguillos et al., 2010; Johnson et al., 2011; Kadkhodaei et al., 2013; Kim et al., 2003; Saijo et al., 2009; Sakurada et al., 1999; Volpicelli et al., 2007). Validation of Nurr1 as a PD therapeutic is primarily derived from mouse models and human data. Homozygous mice lacking Nurr1 fail to generate midbrain dopaminergic neurons and die shortly after birth, heterozygous mice have motor impairments analogous to Parkinsonian deficits, and conditional ablation of Nurr1 in adult animals recapitulates early features of PD with progressive dopaminergic neuropathology (Jiang et al., 2005; Kadkhodaei et al., 2013; Kadkhodaei et al., 2009; Zetterstrom et al., 1997; Zhang et al., 2012a). In patients with PD, the expression of Nurr1 is reduced compared to age-matched controls (Chu et al., 2006; Le et al., 2008; Montarolo et al., 2016; Moran et al., 2007), though only a few, rare polymorphisms in Nurr1 appear to be associated with the disease (Grimes et al., 2006; Le et al., 2003). Stimulation of Nurr1 activity may combat both the reduced dopamine levels and the increased oxidative stress associated with PD.

**Figure 1.**
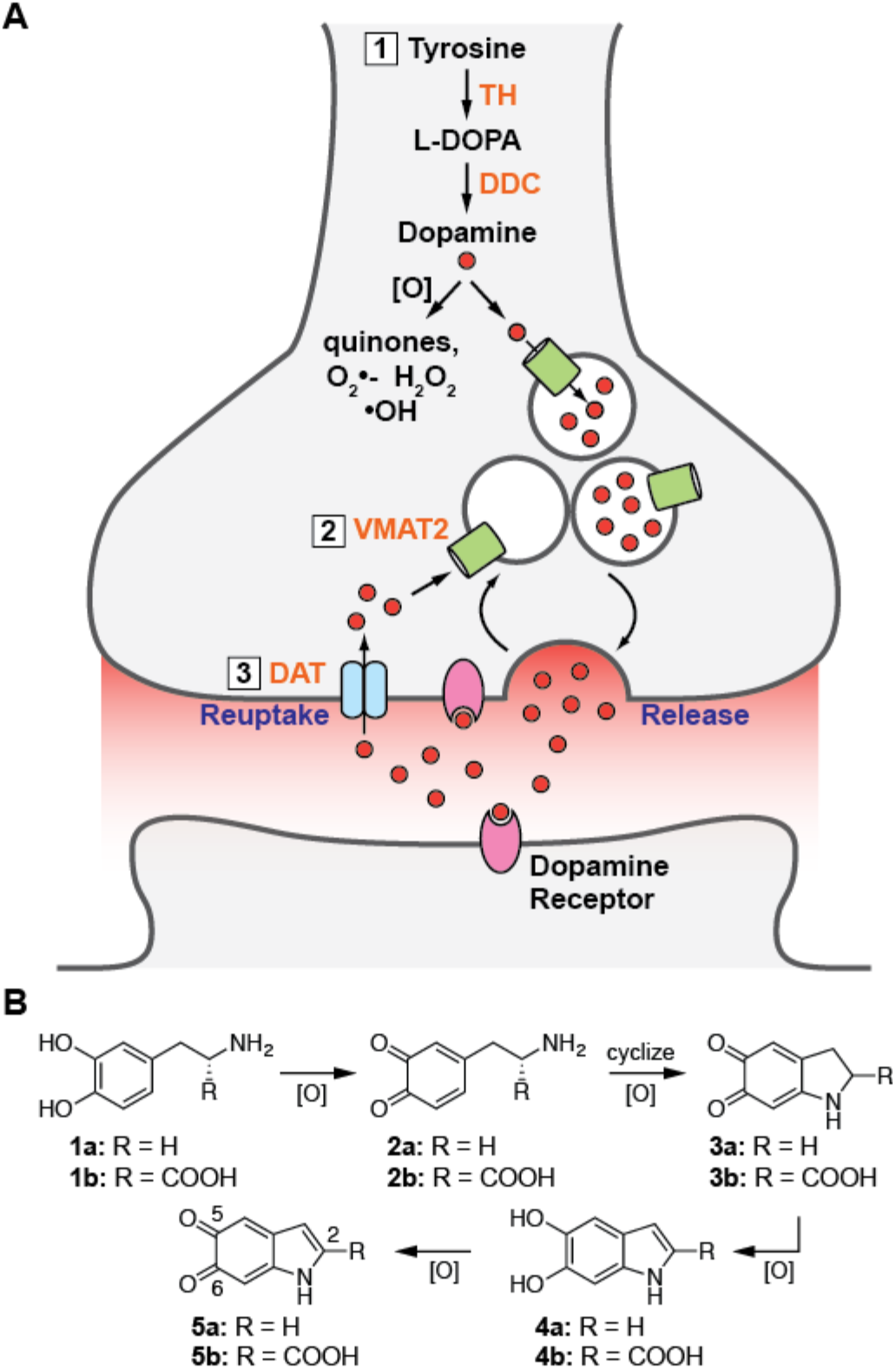
The production and processing of dopamine in neurons. **(A)** Nurr1 controls the dopaminergic phenotype, regulating the genes required for the (1) synthesis (TH, DDC), (2) packaging (VMAT2), and (3) reuptake (DAT) of dopamine. **(B)** Dopamine and L-DOPA auto-oxidize to form reactive indoles and other reactive oxygen species. Only the analogs 1a, 1b, 4a, and 4b are commercially available. 1a = dopamine, 1b = L-DOPA; 2a = dopamine-o-quinone, 2b = dopaquinone; 3a = dopaminochrome, 3b = DOPAchrome; 4a = 5,6-dihydroxyindole (DHI), 4b = 5,6- dihydroxyindole-2-carboxylic acid (DHICA); 5a = 5,6-indolequinone (IQ), 5b = 5,6-dihydroxyindolequinone-2-carboxylic acid (IQCA).

Efforts to identify Nurr1 agonists have been hampered by major gaps in our understanding of the receptor’s structure and regulation. In particular, the only reported crystal structure of the receptor (apo Nurr1), published over 15 years ago, reveals the canonical nuclear receptor (NR) ligand binding pocket is filled by bulky amino acid side chains (Wang et al., 2003). Subsequent efforts to identify ligand binding sites within Nurr1, utilizing NMR studies of the isolated ligand binding domain (LBD), have suggested that small molecules may bind to the receptor in regions corresponding to both canonical and non-canonical ligand binding pockets (de Vera et al., 2016; Kim et al., 2015; Poppe et al., 2007). Phenotypic assays have identified a small number of synthetic ligands that reportedly up-regulate transcription and protein levels of Nurr1 target genes, provide some degree of neuroprotection, and improve behavioral deficits in mouse models (Dong et al., 2016; Kim et al., 2015; McFarland et al., 2013; Smith et al., 2015; Zhang et al., 2012b). However, there is little evidence that any of these ligands directly activate endogenous Nurr1, with the exception of the antimalarial drug amodiaquine (Kim et al., 2015). Endogeneous ligands for Nurr1 have yet to be reported, further limiting our understanding of how this receptor is regulated. Efforts to drug Nurr1 indirectly by targeting the RXR ligand binding domain in Nurr1:RXR heterodimers have demonstrated enhanced expression of Nurr1 target genes by RXR agonists (McFarland et al., 2013; Spathis et al., 2017; Volakakis et al., 2015). This approach may, however, be limited by the established promiscuous association of RXR with other NRs (e.g. RAR, VDR, TR, PPAR, LXR, FXR), and complicated by the apparent repression of Nurr1 transcriptional activity upon complexation with RXR (Perez et al., 2012; Perlmann and Jansson, 1995). The receptor also lacks the canonical co-regulator binding groove, though some reports suggest alternative interaction surfaces for regulatory proteins may be present on the Nurr1 LBD (Codina et al., 2004; de Vera et al., 2016; Volakakis et al., 2006).

Against this backdrop, we set out to identify synthetic and endogenous ligands that bind directly to and regulate the activity of Nurr1. Owing to the pivotal role Nurr1 plays in producing and processing dopamine, and the need for neurons to tightly regulate dopamine levels, we postulated that the receptor might be regulated by dopamine, its biosynthetic precursor L-DOPA, or metabolites of these molecules (Figure 1B). Outside of an acidic environment (e.g. synaptic vesicle), dopamine is unstable (Segura-Aguilar et al., 2014). Following release into the synaptic cleft, excess dopamine is rapidly taken back up into the nerve terminal, traveling through the dopamine transporter, and then repacked into synaptic vesicles, via the vesicular monoamine transporter. Dopamine that is not processed in this way is typically either enzymatically (COMT, MAO) converted to inactive (but oxidatively unstable) metabolites or auto-oxidized into reactive species, including 5,6-dihydroxyindole (DHI) and 5,6-indolequinone (IQ) (Figure 1B), which are the focus of this work (Meiser et al., 2013). The reactive compounds polymerize to form neuromelanin, a chromogenic pigment of uncertain function that accumulates in and stains midbrain dopaminergic neurons black in healthy individuals (Fedorow et al., 2006; Zucca et al., 2017), or are otherwise quenched by direct conjugation to scavenging small molecules (e.g. glutathione) or protein thiols (Sulzer and Zecca, 2000). Using biophysical, structural, and biological assays, we evaluated the interaction of Nurr1 with oxidative metabolites of L-DOPA and dopamine. We found that DHI binds directly to Nurr1 in a non-canonical ligand binding pocket, forming a covalent adduct by reacting as the indolequinone (IQ) with an endogenous cysteine residue (Cys566). In both cultured cells and zebrafish, DHI stimulates Nurr1 transcription, including upregulating the target gene underlying the management of excess cytoplasmic dopamine (i.e. VMAT2).

## RESULTS

### DHI Binds Directly to the Nurr1 Ligand Binding Domain

Differential scanning fluorimetry (DSF) is a fluorescence-based thermal shift assay frequently used to identify specific interactions between small molecules and proteins (Niesen et al., 2007); compounds that bind directly to and stabilize a protein typically increase the melting temperature (Tm). Using DSF, we found that DHI produces a concentration-dependent increase (1°C at 50 μM) in the Tm of the Nurr1 ligand binding domain (LBD) (Figure 2A, C), whereas even the structurally related C2-substituted metabolite 5,6-dihydroxyindole carboxylic acid (DHICA) does not alter the Tm (Figure 2B, D). DHI binding is detected at concentrations above 5-10 μM and there is some evidence of site saturation at the highest concentration tested (50 μM); it was not possible to obtain data at higher concentrations due to apparent polymerization of DHI, which is accelerated upon heating. We also observed that DHI, but not DHICA, produces a concentration-dependent decrease in the overall fluorescence, possibly due to precipitation of the protein caused by DHI polymers (Supplemental Figure 1).

**Figure 2.**
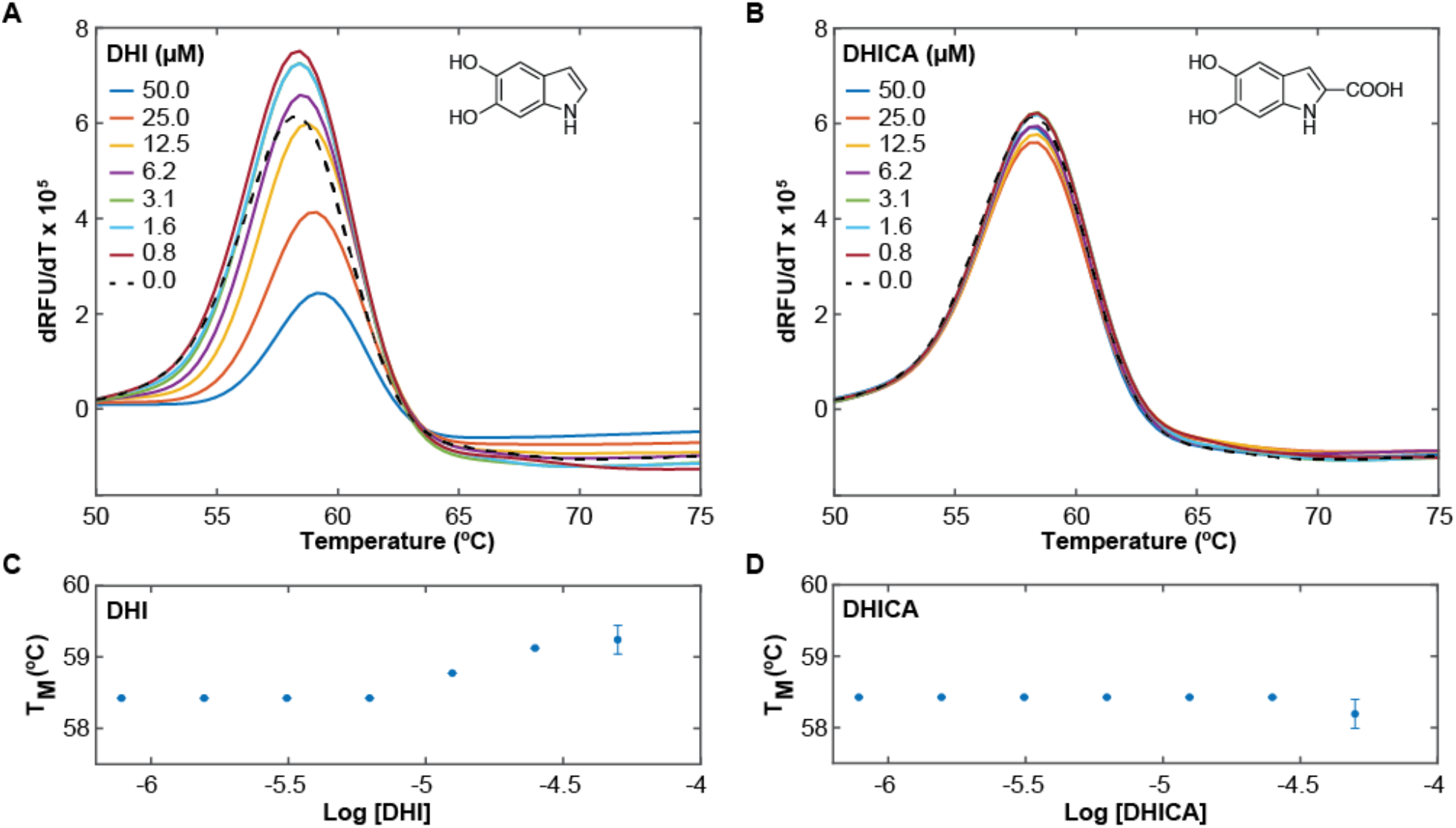
DHI but not DHICA thermodynamically stabilizes the Nurr1 LBD. DSF assays were carried out on the Nurr1 LBD (4 μM) in the presence of the dye SYPRO orange (2.5x) and varying concentrations of DHI or DHICA. DHI, but not DHICA, produces a concentration-dependent increase in the Tm. **(A, B)** Plot of the temperature-derivative of the fluorescence signal versus temperature (dRFU/dT vs temperature), for (A) DHI and (B) DHICA, where the maximum (peak) of the derivative plot identifies the Tm. **(C, D)** Plot of the change in Tm as a function of the concentration of (C) DHI and (D) DHICA. Experiments were performed in triplicate. Error bars are standard deviations from n = 3.

We confirmed direct binding of DHI to the Nurr1 LBD using surface plasmon resonance (SPR). Biotinylated Nurr1 LBD was immobilized on a biotin-capturing chip and screened against increasing concentrations of mobile analytes. As above, DHI (but not DHICA, dopamine, or L-DOPA; Supplemental Figure 2), binds at micromolar concentrations directly to Nurr1 in a concentration-dependent manner and with kinetic features suggestive of a covalent interaction (Figure 3A). In particular, rather than exhibiting step-like ‘fast-on, fast-off’ kinetics typical of reversible small molecule (< 300 Da) binding, DHI appears to not dissociate from the receptor on the timescale of minutes, under these conditions.

**Figure 3.**
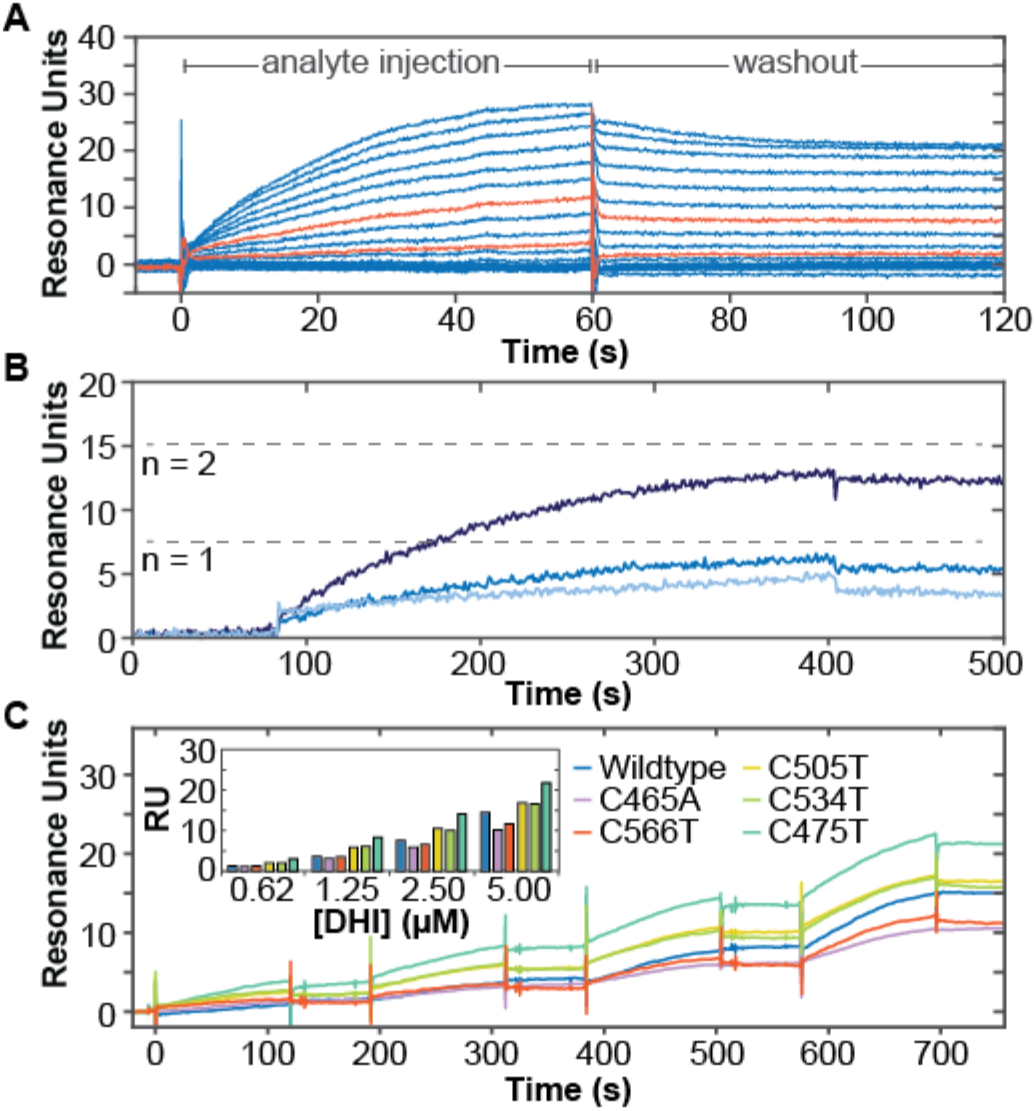
Direct association of DHI with the Nurr1 LBD measured by surface plasmon resonance. Biotinylated Nurr1 LBD was coupled to a streptavidin-coated biosensor chip and exposed to DHI at varying concentrations, either without (A, C) or with (B) surface regeneration between DHI exposures. **(A)** Concentration-dependent association of DHI with Nurr1 LBD. DHI was injected at 15 increasing concentrations (16.8 μM DHI, 0.6x dilutions down to 0.01 μM; 60 s association time). DHI injections at 0.17 and 0.78 μM are highlighted in orange. **(B)** Concentration-dependent association of DHI with Nurr1 LBD (0.025, 0.25, 2.5 μM DHI; 320 s association time), with surface regeneration between DHI injections. Dashed lines indicate theoretical Rmax for a 1:1 (n =1) or 1:2 (n = 2) DHI:Nurr1 ratio. **(C)** Concentration- and time-dependent association of DHI with Nurr1 wildtype and each of five cysteine mutants. DHI was injected at four increasing concentrations (0.62, 1.25, 2.50, 5.00 μM; 120 s association time, 60 s dissociation time). Raw responses were normalized to the initial protein captured. **(Inset)** Bar graph comparing plateau responses for each protein at the four DHI concentrations tested.

To better characterize the kinetics of the DHI-Nurr1 interaction, we performed an additional SPR experiment in which the surface of the chip was regenerated (replacing the Nurr1 LBD) after each exposure to DHI to ensure that each response reflects DHI binding to naïve apo Nurr1 LBD. (Figure 3B). These experiments confirm that DHI exhibits the features of a specific, covalent binding event to Nurr1 by showing (i) a concentration-dependent increase in association rate, (ii) convergence of the association towards steady state at the highest concentration tested (2.5 μM; 320 s contact time), (iii) maximal association responses consistent with the theoretical maximal response (Rmax^1^) for one or possibly two molecules of DHI binding to each Nurr1 LBD, ruling out super-stoichiometric, non-specific binding.

As DHI and DHICA have been shown to form covalent adducts with cysteine, including in the process of pheomelanin formation (d’Ischia et al., 2005; Dischia et al., 1987; Mason and Peterson, 1965), we hypothesized that DHI might bind covalently to the Nurr1 LBD through one of its five conserved cysteine residues (Supplemental Table 1). Consistent with this hypothesis, SPR experiments show that DHI binding to Nurr1 is reduced in the Cys566Thr and Cys465Ala mutants, but not in Cys505Thr, Cys534Thr, or Cys475 mutants (Figure 3C). Neither the Cys566Thr nor the Cys465Ala mutation eliminated DHI binding, suggesting that DHI binds to both Cys566 and Cys465. Examples of other electrophilic nuclear receptor ligands that also react with more than one cysteine residue have been previously reported (Reese et al., 1992). Unfortunately, we were not able to detect the DHI modification by mass spectroscopy, consistent with previous reports showing that “dopamine-modified” proteins (i.e. proteins exposed to dopamine under oxidizing conditions) could not be detected by mass spectroscopy (Armarego and Waring, 1983; Belluzzi et al., 2012; Bisaglia et al., 2007; Girotto et al., 2012; Hastings, 2009; Kuhn et al., 1999; LaVoie et al., 2005; Whitehead et al., 2001). It is possible that the ionization process drives the neutral loss of these adducts and/or that the adducts are chemically reversible. Indeed, the reversibility of protein thiol Michael additions to drugs with enone systems has been previously observed (Lee et al., 2002). Unfortunately, Nurr1 LBD with four or five of the five cysteine residues mutated could not be expressed and/or purified, making it difficult to unambiguously determine the site or sites of DHI binding in this manner. As discussed below, we confirmed specific binding of DHI to Cys566 by x-ray crystallography, but cannot rule out binding to a secondary site, most likely Cys465. Notably, Cys465 is oxidized in one of the three chains in the x-ray structure (Supplemental Figure 3), the biological significance of which is worthy of further exploration.

### The Quinone Form of DHI Binds to the Nurr1 LBD within a Non-Canonical Ligand Binding Pocket

We characterized the structure of the Nurr1 LBD co-crystalized with DHI at the atomic scale using protein crystallography. The structure of the complex, determined at 3.2 Å resolution (PDB: 6DDA; Supplemental Table 2), reveals that the metabolite binding site sits adjacent to the canonical ligand binding pocket^2^, and confirms the modification of Cys566 (Figure 4). Structural alignment of the peptide backbones for liganded and apo Nurr1 LBD (pdb: 1OVL) reveals an overall C*α* RMSD of 0.57 Å and highlights two important differences between the structures (Figure 4A). First, Cys566 appears in two distinct rotameric conformations in the structures; in the apo structure the thiol points towards the vestigial canonical ligand binding pocket, whereas in the liganded structure it points towards a pocket formed by H11, H12, and the bend in the middle of H4/5. Second, a 1.5 Å positional shift is observed between the two structures at the end of H12, a classic regulatory element among NRs. Otherwise, the two structures are very similar.

**Figure 4.**
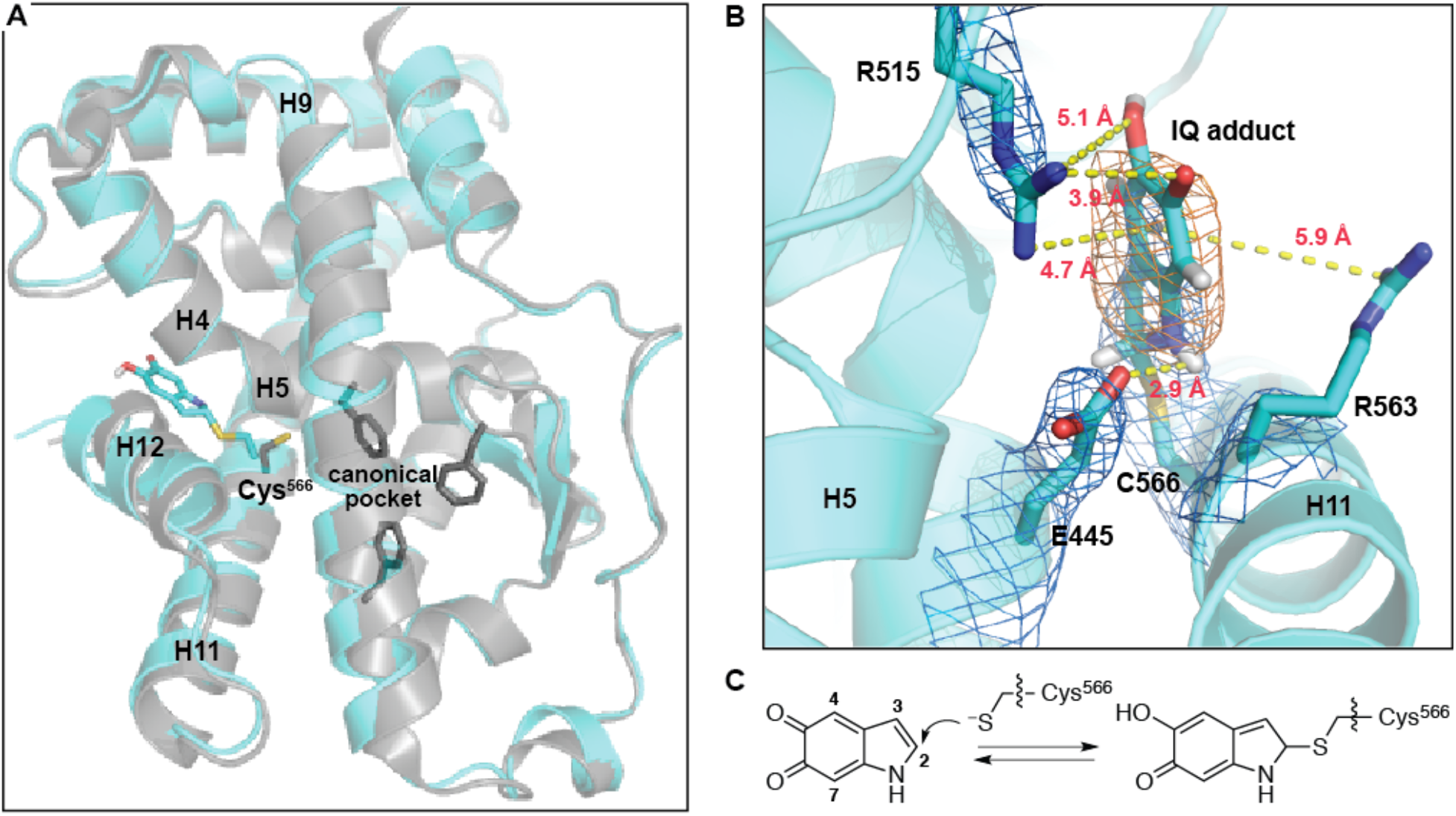
Crystal structure of 5,6-indolequinone bound to the Nurr1 LBD. **(A)** Structural alignment of the IQ-bound structure (cyan; PDB: 6DDA, chain B) with the apo structure (grey, PDB: 1OVL, chain B) using H1 and H7 peptide backbone atoms for alignment. The C-alpha RMSD calculated across the entire ligand binding domain using this alignment is 1.16 Å. Shown as sticks: Cys566 IQ adduct (cyan), Cys566 from the apo structure (grey), residues in the apo structure that fill the canonical ligand binding pocket (Phe447, Phe464, Phe479; grey). **(B)** IQ forms a covalent adduct with Cys566. Depicted adduct corresponds to IQ linked at the C2 atom to Nurr1. Electron density map (2mFo-DFc, contoured to 1σ) indicated with blue mesh. Polder difference map (orange mesh) is contoured to 4.5σ in strength. Polar interactions between the Cys566 IQ adduct and Glu445 and Arg515 are highlighted; dashed yellow lines indicate distances between non-hydrogen atoms. Potential cation-pi interactions between the ligand and Arg515 and/or Arg563 are also indicated with dashed yellow lines. Weak electron density for Arg563 suggests it is dynamic. **(C)** Chemical reaction between Cys566 and IQ leading to adduct formation.

To enhance the signal of the ligand electron density, we generated Polder difference maps, which take into consideration bulk-solvent flattening effects that traditional OMIT maps do not (Figure 4B, Supplemental Table 3). The ligand density is most consistent with a covalent bond between the Cys566 sulfur atom and C2 atom of IQ, the auto-oxidation product of DHI (Figure 4C); this density is not observed surrounding any of the other cysteine residues in the protein (Supplemental Figure 3) Modeling with IQ attached via the C3, C4 or C7 atoms, which have previously been shown to be reactive sites (d’Ischia et al., 2011; d’Ischia et al., 1987), not only fit the ligand density more poorly, but also generated unavoidable steric clashes with protein side chains in the pocket.

Since it is not possible to unequivocally conclude from the electron density map that the metabolite is bound as the quinone, we employed quantum mechanical calculations to characterize the transition states and energy barriers for thiolate attack with both redox states (DHI and IQ) at each of the possible electrophilic sites (C2, C3, C4, C7). These studies revealed that the IQ C2-adduct is the most likely reaction product. Reaction between the thiolate and quinone species (IQ) at positions 2, 4 and 7 are all predicted to generate stable adducts, with transition state energy barriers of 6.3, 14.7 and 16.7 kcal/mol, respectively, relative to the reactants, whereas reaction at C3 is predicted to be unfavorable as the energies for the transition state and the corresponding product are both high (E_act_ > 30.0 kcal/mol) (Supplemental Figure 4). Interestingly, the barrier to dissociation of the C2-quinone adduct is only ~3 kcal/mol, suggesting that the covalent adduct could be reversible under some conditions. Reaction between the thiolate and the catechol species (DHI) is unlikely, as all four adducts are predicted to be high-energy, unstable states that would spontaneously dissociate (Supplemental Figure 5).

Modeling of the IQ C2-adduct into the electron density reveals that several intermolecular interactions are expected to stabilize the binding of DHI/IQ, and the covalent IQ-protein complex, including: i) hydrogen bonds between the guanidinium of Arg515 (H8-9 loop) and the C5 and C6 oxygens of DHI and IQ; ii) a hydrogen bond between the carboxylate of E445 (H4/5) and the NH of DHI and IQ; iii) a cation-pi interaction between Arg515 and/or Arg563 (H11) and the pi systems of DHI and IQ^3^ (Figure 4B). One or both of these arginine residues may also stabilize formation of the oxyanion resulting from nucleophilic attack of the thiol. In the apo Nurr1 LBD structure the DHI/IQ binding site is occupied by ordered water molecules (two-four, depending on the chain); the release of these waters back into the bulk solvent (i.e. receptor desolvation) is entropically favorable and may also contribute to the initial non-covalent binding event. These interactions may also explain the preferential binding of DHI over DHICA; the latter contains a C2-carboxylate that renders it too big to fit within the metabolite pocket. An unresolved question in this work is whether DHI first autooxidizes to IQ in solution, after binding non-covalently to Nurr1 near Cys566, or both; the extreme instability of IQ makes it challenging to study. As noted, many of the favorable interactions observed in the IQ-Nurr1 adduct crystal structure would also stabilize non-covalent interactions with DHI, and we speculate that the positive charges of Arg515 and Arg563 might decrease the pKa of DHI hydroxyl groups, increasing its rate of oxidation to IQ, which generates two protons.

### DHI Stimulates Nurr1 Activity

To probe the effect of DHI on the activity of Nurr1 in cultured cells, we utilized a reporter assay in which the NR LBD is fused to the Gal4 DBD, and binding of the chimeric protein to a reporter >plasmid sequence (UAS cis-element) drives the expression of firefly luciferase (Castro et al., 1999). For these assays, JEG3 cells were co-transfected with an expression plasmid encoding the wildtype Nurr1 LBD fused to the Gal4 DBD (pM-Nurr1-Gal4) or the Gal4 DBD alone (pM-Gal4), along with a reporter plasmid (pGL4.35) encoding the firefly luciferase gene driven by an upstream 9xGal4, and an internal control plasmid (pRL-TK) expressing renilla luciferase. Transfected cells were incubated with DHI, DHICA, or DMSO control for 6 h before luciferase signals were measured. In these assays, DHI produces a concentration-dependent increase in firefly luciferase expression, with a statistically significant effect at 10 μM DHI, and a 1.6-fold over basal increase in activation at 100 μM DHI (Figure 5). Basal activation is seen in the absence of DHI and DHICA (i.e. DMSO only), in agreement with the constitutive Nurr1 activation reported by others (Castro et al., 1999; Wang et al., 2003). Concentrations of DHI above 100 μM were cytotoxic, preventing us from obtaining an EC_50_.

**Figure 5.**
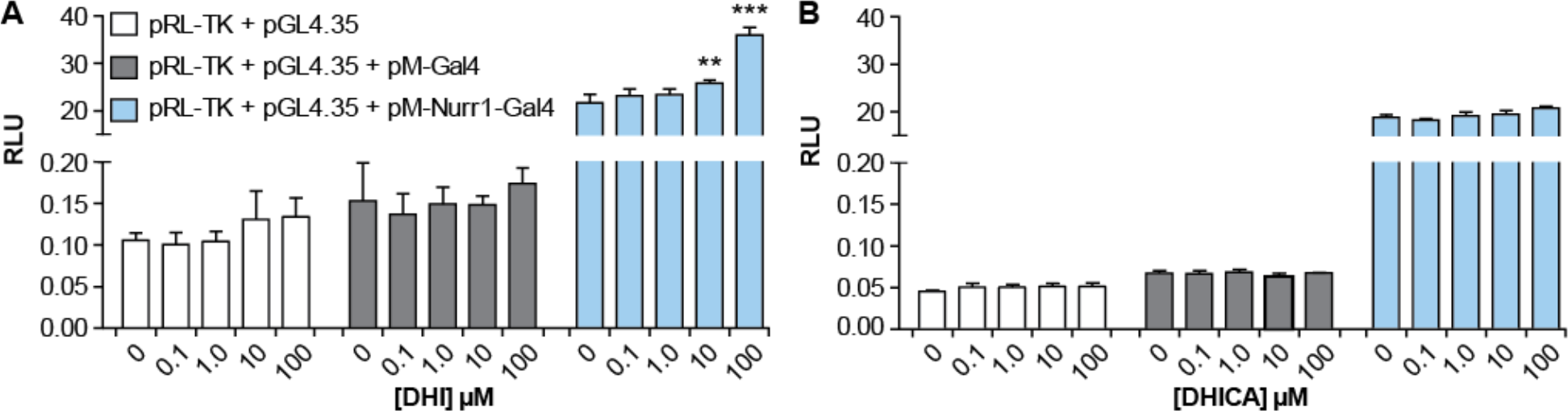
DHI stimulates Nurr1 mediated transcription. JEG3 cells were co-transfected with an internal control plasmid (pRL-TK) expressing renilla luciferase, a reporter plasmid (pGL4.35) encoding the firefly luciferase gene driven by an upstream 9xGal4 element, and an expression plasmid encoding the Nurr1 LBD fused to the Gal4 DBD (pM-Nurr1-Gal4) or the Gal4 DBD alone (pM-Gal4). Cells were incubated with varying concentrations of **(A)** DHI or or **(B)** DHICA for 6 h before measuring the luciferase signal (RLU: relative luminometer units). One-way Analysis of Variance (ANOVA), in comparison to the data with 0 μM compound (DMSO only), **p < 0.01, ***p < 0.001.

To study the effect of DHI on the activity of Nurr1 in neurons, we characterized the transcription of several Nurr1 target genes in zebrafish, a model organism that has been used for studying Parkinson’s disease (Xi et al., 2011; Zhang et al., 2017). The zebrafish genome encodes two isoforms of Nurr1, the NR4A2A isoform expressed predominantly in dopaminergic neurons and the NR4A2B isoform expressed predominantly in the eye (Blin et al., 2008). The LBD of the neuronal isoform has remarkably high sequence identity, 95%, with the human Nurr1 LBD, and all of the cysteine residues are conserved (Supplemental Table 1). In these assays, zebrafish embryos three days post-fertilization (3 dpf) were exposed to DHI for either 6 h or 24 h, after which RNA was extracted and subjected to RT-qPCR analysis (Figure 6). At 6 h post treatment with 100 μM DHI, only the transcript levels of genes coding for the transporters of dopamine (*vmat2*, *dat*) show statistically significant increases (Figure 6A). At 24 h post-treatment, the increased expression of *vmat2* remains significant, and there are also statistically significant increases in the transcript levels for the rate-limiting biosynthetic enzyme tyrosine hydroxylase (*th*) (Figure 6B).

**Figure 6.**
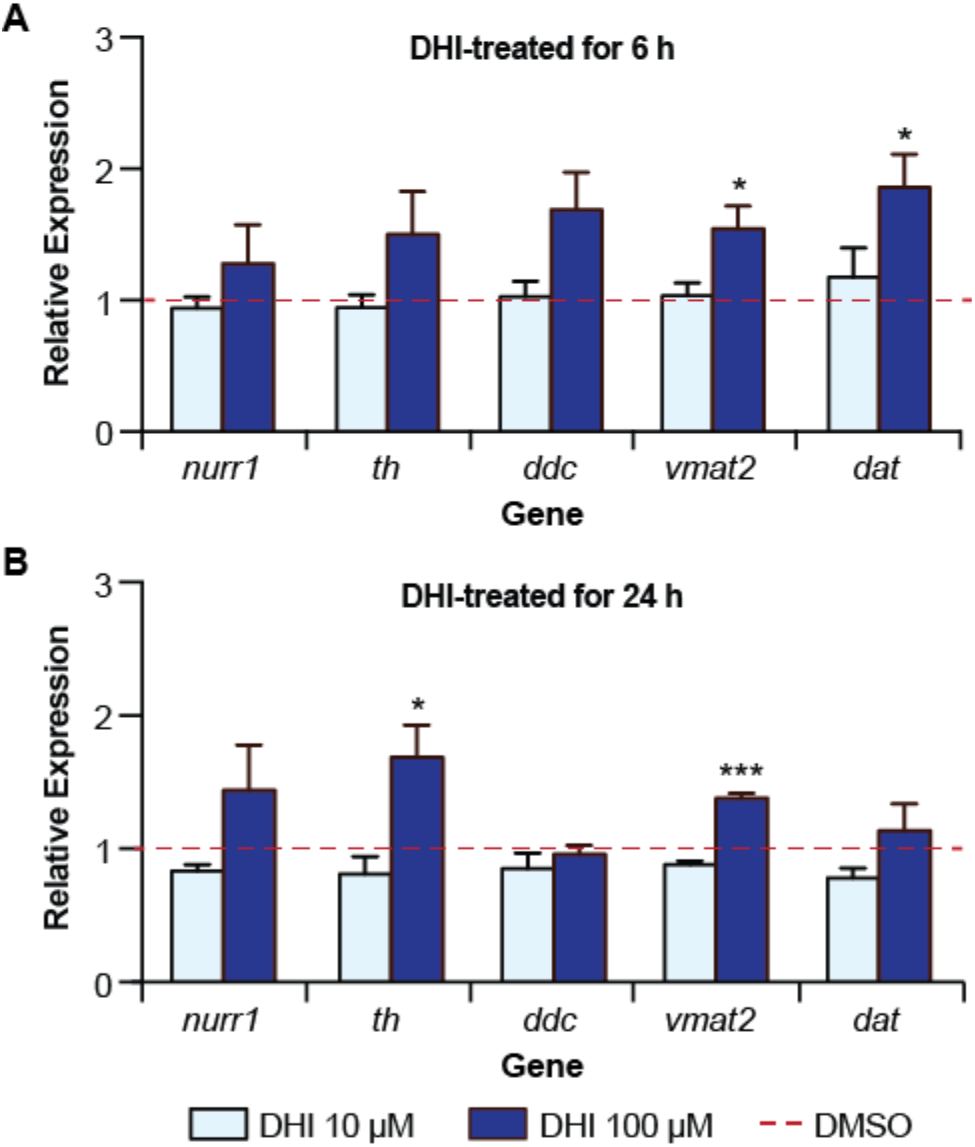
DHI drives the transcription of a subset of Nurr1-dependent genes in zebrafish. **(A)** qPCR analysis of mRNA from wildtype zebrafish larvae 78 hpf following 6 h treatment with DHI (100 μM) show statistically significant increases in transcripts of *vmat2* and *dat* compared to DMSO control (red bar). **(B)** qPCR analysis of mRNA from wildtype zebrafish larvae 96 hpf after 24 h of DHI (100 μM) treatment show statistically significant increases in transcripts of *th* and *vmat2* compared to DMSO control (red bar). Transcript levels for target genes were normalized to the housekeeping gene *elf1a* and fold change was compared to gene expression levels from DMSO-treated larvae. Results are from four independent experiments. Relative average expression ± SEM; *p < 0.05, **p < 0.01, ***p < 0.001 by Student’s t-test.

## DISCUSSION

Considerable evidence suggests that dysregulation of dopamine is both a contributor to and consequence of PD (Burbulla et al., 2017; Hastings, 2009; Jenner, 2003; Lotharius and Brundin, 2002; Sulzer et al., 2000). Metabolism of dopamine produces reactive oxygen species (ROS) and quinones, and the formation of these toxins is exacerbated by excessive levels of cytoplasmic dopamine (VMAT2 dysfunction), increased levels of ROS (mitochondrial dysfunction), and other forms of oxidative stress—all conditions associated with PD. The transcriptional regulator Nurr1 plays a pivotal role in maintaining dopamine homeostasis, regulating the synthesis, packaging, and re-uptake of the neurotransmitter. The regulation of Nurr1 itself is incompletely understood, however, partly owing to the absence of a well-defined ligand binding pocket within the receptor. Delineating a binding site for small molecules within Nurr1 is a critical step toward understanding this receptor’s role in and potential effect on PD.

In this study, we used biophysical and structural assays to identify a binding site for a specific dopamine metabolite within the Nurr1 LBD. We found that 5,6-dihyroxyindole (DHI) binds to Nurr1 within a non-canonical binding site, forming a covalent adduct as the indolequinone (IQ) with Cys566. The interaction with DHI/IQ is detectable by surface plasmon resonance in the high nanomolar range (>0.25 μM), a concentration consistent with sensing cytoplasmic dopamine under conditions of oxidative stress (Eisenhofer et al., 2004; Mosharov et al., 2003; Mosharov et al., 2006; Olefirowicz and Ewing, 1990; Omiatek et al., 2013; Pifl et al., 2014), and starts to show evidence of saturable binding at low micromolar concentrations (2.5 μM, Figure 3). In functional assays, DHI stimulates Nurr1 activity, driving the transcription of genes controlling dopamine homeostasis. These data suggest that Nurr1 functions as a sensor for oxidative stress in dopaminergic neurons, responding directly to a specific oxidative metabolite of dopamine.

### DHI Binds to Nurr1 within a Regulatory Hotspot

Nuclear receptors are highly modular structures, typically with multiple surfaces engaging in protein-protein interactions and accessibility to these sites allosterically regulated by the binding of small molecules within the LBD (Gallastegui et al., 2015; Gronemeyer et al., 2004; Huang et al., 2010; Kumar and McEwan, 2012). Consistent with this established view of NR regulation, the DHI/IQ binding site is situated among several regulatory surfaces and metabolite binding produces a structural change in H12, a key regulator of NR function (Figure 7). In particular, metabolite binding produces 1.5 Å shift in H12 that potentially impacts the binding of co-regulator proteins (Codina et al., 2004), nuclear export of the receptor (Garcia-Yague et al., 2013), homodimerization with Nurr1 (Zhan et al., 2012), and heterodimerization with RXR (Aarnisalo et al., 2002).

**Figure 7.**
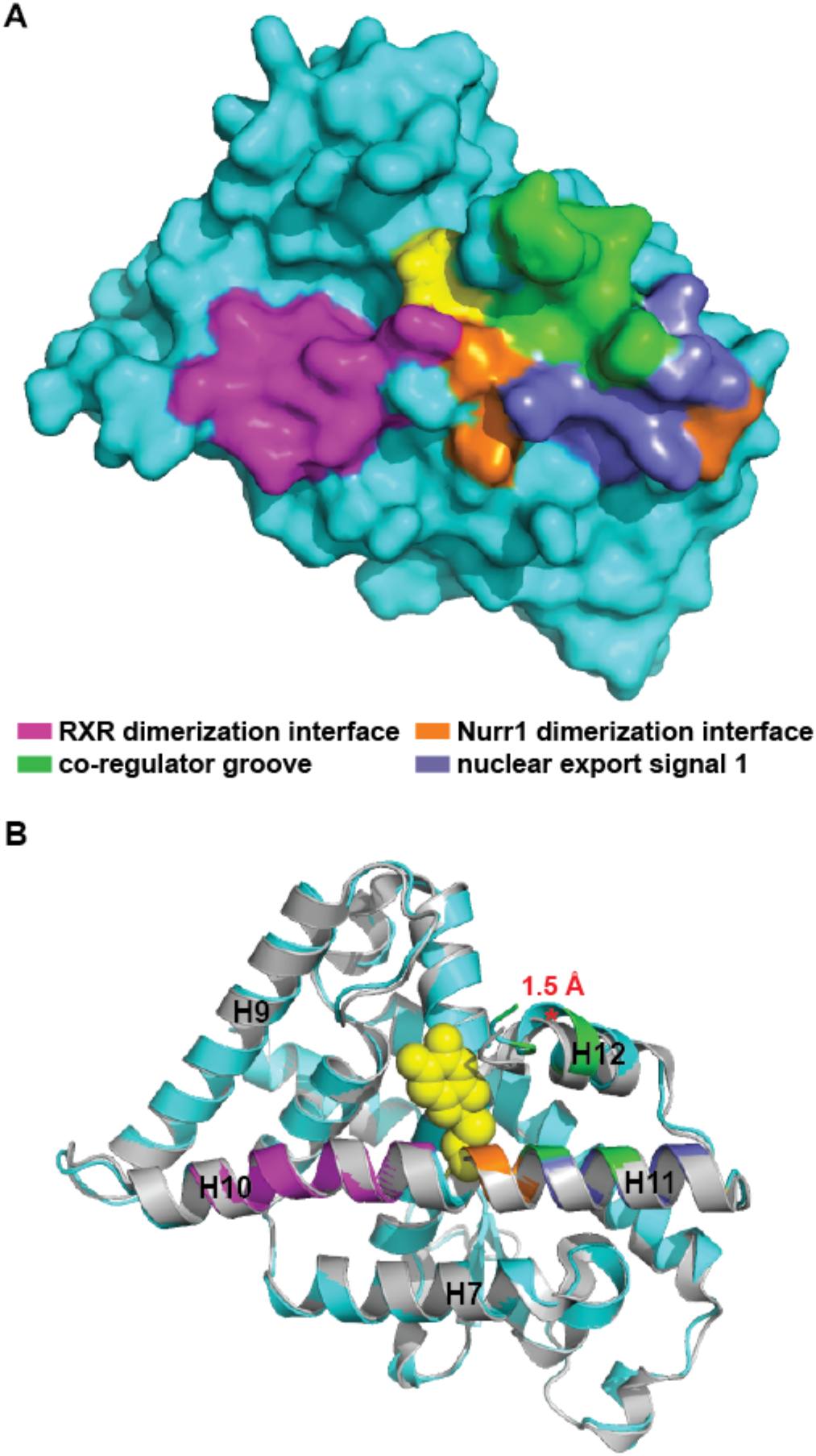
The DHI/IQ binding site is situated between key regulatory elements of Nurr1 function. Cys566-IQ adduct shown as yellow spheres. **(A)** Surface rendering of the Nurr1-IQ structure highlighting the relationship between the bound metabolite and the RXR heterodimerization interface (magenta), Nurr1 homodimerization interface (orange), co-regulator groove (green), and nuclear export signal 2 (slate). **(B)** Superposition of the liganded Nurr1 structure (cyan) with the apo structure (grey), highlighting the shift (red asterisk*) between alpha-carbon residues at the end of H12 in the two structures. Nurr1 poses are identical in (A) and (B). Alignment is identical to that described for Figure 4a.

The conformational changes affected by DHI/IQ appear to alter the activity of Nurr1, upregulating the expression of Nurr1 target genes. Nurr1 reportedly signals as a monomer, homodimer, and heterodimer in neurons, with each conformation recognizing distinct DNA response elements and thus regulating transcription of different target genes (Maira et al., 1999; Paulsen et al., 1995; Perlmann and Jansson, 1995; Philips et al., 1997; Volpicelli et al., 2012; Zetterstrom et al., 1996). Our data suggests that DHI/IQ binding to Nurr1 biases the oligomerization state of the receptor toward the monomer, as two of the genes upregulated by the metabolite (*vmat2* and *th*) both contain promoter sequences that are recognized by full-length Nurr1 monomers (i.e. NBRE and NBRE-like response elements).^4^

### Nurr1 Senses and Responds to Oxidative Stress

A number of studies have shown that auto-oxidized dopamine metabolites can covalently modify proteins intimately involved in PD, such as SOD2, TH, DJ1, DAT, DHPR, alpha-synuclein, and parkin (Armarego and Waring, 1983; Belluzzi et al., 2012; Bisaglia et al., 2007; Girotto et al., 2012; Hastings, 2009; Kuhn et al., 1999; LaVoie et al., 2005; Whitehead et al., 2001). In all of these cases, the modifications inhibit normal protein function, often by driving the formation of higher order protein species (e.g. crosslinked protein) and/or by reducing protein stability.^5^ In contrast, here we provide the first example of a specific protein modification by a dopamine metabolite that stimulates function.

We observed that acute (6 h) treatment of zebrafish with DHI stimulates the transcription of genes underlying the reuptake and packing of dopamine (*dat*, *vmat2*), whereas prolonged (24 h) treatment stimulates transcription of genes underlying the synthesis and packaging genes (*th*, *vmat2*). The observed upregulation of VMAT2 gene transcripts is particularly interesting as the functioning of this vesicular transporter is an important determinant of dopamine-related oxidative stress, neurodegeneration, and PD (Alter et al., 2013; Guillot and Miller, 2009). For example, reduced vesicular storage of dopamine has been shown to result in progressive neurodegeneration in mouse models (Caudle et al., 2007; Chen et al., 2008; Ulusoy et al., 2012). Furthermore, defects in VMAT2 function are observed in synaptic vesicles isolated from the striatum of patients with PD (Pifl et al., 2014). Conversely, a gain of function VMAT2 haplotype in humans has been shown to be protective in PD (Brighina et al., 2013; Glatt et al., 2006) and upregulation of VMAT2 in mouse models of PD has been shown to provide protection against neurodegeneration (Lohr et al., 2014). Finally, drugs that inhibit VMAT2 function or otherwise increase levels of cytoplasmic dopamine (e.g. methamphetamine, cocaine) are associated with degeneration of dopaminergic neurons and increased risk for PD (Guillot et al., 2008; Krasnova and Cadet, 2009). Thus, in addition to its fundamental role in neurotransmission, VMAT2 also plays a neuroprotective role by sequestering dopamine in acidic synaptic vesicles.

### Nurr1 is a Ligand-inducible Transcription Factor

Although Nurr1 is widely recognized as a therapeutic target for PD, the absence of a well-defined ligand binding pocket has hampered efforts to capitalize on this potential. Previous reports have hinted at canonical and non-canonical ligand binding sites within the Nurr1 LBD (Supplemental Figure 6) (de Vera et al., 2016; Kim et al., 2015; Poppe et al., 2007). The binding site for DHI/IQ is most closely approximated by Poppe et al., who highlighted the proximity of Cys566 to a binding site for benzimidazole-containing compounds. Additional x-ray structures of liganded Nurr1 further underscore the existence of multiple small molecule binding sites within the LBD (manuscript in preparation).

Our findings demonstrate that Nurr1 is a ligand-inducible transcription factor. While DHI itself is an unstable molecule, auto-oxidizing and polymerizing with itself and other dopamine metabolites, the indole core is a tractable scaffold for developing stable compounds targeting Nurr1. Further, the reactivity of Cys566 suggests avenues for developing both reversible and irreversible covalent compounds targeting the receptor. Such compounds could impact the progression and symptoms of PD. Improving the packaging of dopamine into vesicles by upregulating transcription of VMAT2 may slow the progression of Parkinson’s disease, and extend the treatment window for L-DOPA by reducing side-effects. Increasing the synthesis of dopamine by upregulating transcription of the rate limiting biosynthetic enzyme, TH, offers an orthogonal method for increasing dopamine levels.

## SIGNIFICANCE

Dysregulation of dopamine homeostasis contributes to Parkinson’s disease (PD). The nuclear receptor Nurr1 plays a central role in dopamine homeostasis, regulating the transcription of genes governing the synthesis, packaging, and reuptake of dopamine. Efforts to capitalize on Nurr1’s potential as a therapeutic target for PD have been hampered by the absence of a defined ligand-binding pocket within the receptor. We found that the dopamine metabolite 5,6- dihydroxyindole (DHI) binds to Nurr1 within a non-canonical pocket, forming a covalent adduct as the 5,6-indolequinone (IQ) with Cys566. In functional assays, DHI stimulates Nurr1 activity, upregulating the transcription of several genes, including the vesicular monoamine transporter 2. These results suggest that Nurr1 may be regulated by an endogenous metabolite, contradicting previous suggestions that it has no small molecule regulation.

## EXPERIMENTAL PROCEDURES

### Nurr1 LBD Constructs

Nurr1 LBD constructs and sequences are described in detail in Supplemental Table 4. All constructs were subcloned into the pET-DUET Vector (Novagen) within the first multiple cloning site (MCS) and in frame with the N-terminal 6x His tag. The wildtype constructs used for the surface plasmon resonance assays and x-ray crystallography were subcloned into the pET-DUET vector utilizing the BamHI and HindIII restriction sites within the MCS. Note: The N-terminal Ser residue in the crystallography construct (residues Ser-328-598) is an artifact of TEV cleavage (TEV recognition site ENLYFQS, with cleavage between the Q and S) and was not present in the previously reported crystallization construct (Wang et al., 2003). The wildtype construct used for the thermal shift assays was subcloned into the pET-DUET vector using Gibson Cloning. The linearized pET-DUET vector was prepared by PCR amplification, starting at the first multiple cloning site (forward primer: TGCTTAAGTCGAACAGAAAGTAATCG, reverse primer: GGTATATCTCCTTCTTAAAGTTAAAC). The gene inserts (synthesized by Integrated DNA Technologies) including additional 5’ and 3’ Gibson overlap sequences (5’ overlap: GTTTAACTTTAAGAAGGAGATATACC; 3’ overlap: TGCTTAAGTCGAACAGAAAGTAATCG) were then incorporated into the linearized pET-DUET vector using Gibson Assembly (New England BioLabs). The cysteine mutant constructs were prepared using Gibson Cloning as described above (Cys566Thr; gene insert prepared by Integrated DNA technologies) or by custom gene synthesis (Cys465Ala, Cys475Thr, Cys505Thr, Cys534Thr; prepared by GenScript) starting from the wildtype 6xHis Tag – Avi Tag – Nurr1 LBD (363-598) construct (see Supplemental Table 4).

### Protein Expression and Purification

#### Nurr1 LBD Preparation for Crystallography

The pET-DUET plasmid harboring the Nurr1 LBD (residues Ser-328-598; Supplemental Table 4) was transformed into BL21(DE3) (New England BioLabs) or BL21-CodonPlus(DE3)-RIL (Agilent) RILS cells. A single colony was used to inoculate 100 mL of Lysogeny Broth Lennox (LB) medium supplemented with 50 μg/mL ampicillin (as well as 34 μg/mL chloramphenicol in the case of the BL21-CodonPlus strain) and the culture was allowed to grow overnight at 37°C and 180 rpm. A 10 mL aliquot of the overnight culture was used to inoculate 1 L of LB supplemented with antibiotics as above and the resulting culture was allowed to grow at 37°C, 180 rpm until reaching an OD_600_ of 0.6 AU, at which time the incubator and cultures were dropped to 16 °C, 180 rpm. Once the culture reached an OD_600_ of 0.8 AU, protein expression was induced by addition of 250 μL of 1M IPTG per 1 L of culture (final concentration 250 μM IPTG) and allowed to proceed overnight (~16 h) at 16°C and 180 rpm. Cells were then harvested by centrifugation (3500 x g, 20 min), resuspended in ~20 mL of Lysis Buffer per liter of culture using and 18-gauge needle, and lysed by continuous passage at 15,000 psi using a C3 Emulsiflex (Avestin). The soluble Nurr1 LBD was purified from the supernatant using metal affinity chromatography, anion exclusion chromatography, and size exclusion chromatography, as follows. The insoluble fraction was removed by centrifugation (35,000 x g, 30 min). The supernatant was then mixed with 1 mL of TALON resin (Clontech) per liter of culture and allowed to incubate for 2 h at 4°C with continuous inversion on a rotating wheel. The protein-bound resin was then washed sequentially with 100 mL each of Base Buffer supplemented with 20 mM MgCl_2_ and 8 mM ATP, Base Buffer supplemented with 100 mM KCl, and Base Buffer supplemented with 25 mM imidazole, prior to elution with 5 mL of Base Buffer supplemented with 300 mM imidazole per liter of culture. Protein was then dialyzed against 1 L of Base Buffer overnight (4 °C, with mixing) in the presence of TEV protease. Cleavage was monitored by SDS-PAGE using a 4-12% Bis-Tris (NuPage). The cleaved protein was then purified by anion exchange chromatography over a MonoQ 10/300GL (GE Healthcare Life Sciences) column using a linear NaCl gradient between Mono Q Buffer A and Mono Q Buffer B (0-50% Buffer B over 10 column volumes). Peaks containing Nurr1 were pooled and then allowed to incubate with TALON resin at 4°C for 2 h or overnight, with inversion, to remove any residual uncleaved protein. Flow-through from this reverse metal affinity step was then applied to a S75 16/60 SEC column (GE Healthcare Life Sciences) using Base Buffer as the running buffer. The resulting protein was then concentrated and stored at −80°C for further use. Buffer compositions are as follows: Base Buffer: 50mM HEPES, pH 7.4, 300mM NaCl, 10% Glycerol, 250μM TCEP; Lysis Buffer: Base Buffer + 0.05% Tween 20, 1 cOmplete protease inhibitor tablet (EDTA-free)/50 mL; MonoQ Buffer A: 20mM Tris, pH 7.9, 10% Glycerol, 250μM TCEP; MonoQ Buffer B: MonoQ Buffer A + 1M NaCl.

#### Nurr1 LBD Preparation for Thermal Shift Assays

Nurr1 LBD protein was expressed and purified identically to that used for crystallography, except that the TEV cleavage step and reverse immobilized metal affinity chromatography (i.e. incubation with TALON resin) after ion exchange chromatography were omitted. Accordingly, the resulting Nurr1 LBD protein retains an N-terminal 6xHis Tag.

#### Nurr1 LBD Preparation for Surface Plasmon Resonance Assays

Nurr1 LBD protein was expressed and purified identically to that used for crystallography, with the following exceptions. The pET-DUET plasmid harboring the Nurr1 LBD containing an N-terminal BirA biotinylation site (“avi-tag”) was co-transformed along with the BirA biotin ligase plasmid into BL21(DE3) bacterial cells (New England BioLabs), and 10 mL of warm 5 mM biotin in 10 mM bicine (pH 8.3) per liter of culture was added along with the IPTG at the time of induction. Purification of the biotinylated Nurr1 LBD proceeded as above, except the TEV cleavage step and anion exchange chromatography steps were omitted (i.e. only the initial metal affinity and size exclusion chromatography were performed). Accordingly, the resulting Nurr1 LBD protein is biotinylated and retains an N-terminal 6xHis Tag.

### DHI Preparation

5,6-Dihydroxyindole (Combi-Blocks; QB-2748) and 5,6-dihydroxyindole carboxylic Acid (Santa Cruz Biotech; sc-499091) were each prepared as 10 mM and 100 mM stocks in DMSO, aliquoted in small volumes, and stored at −80°C. Stock solutions were prepared rapidly and away from light to minimize oxidation and subsequent polymerization of the compounds. To monitor the stability of these stocks by ^1^H NMR, 100 mM stocks were also prepared in deuterated DMSO at the same time, stored at −80°C, and re-examined by NMR annually; no significant changes in the spectra were observed at 12 and 24 months. Once thawed, stocks were never re-used. Investigators planning to carry out experiments with commercially available DHI should verify that the distributor stores their stock under inert gas, away from light, and ideally at < 25°C. Combi-Blocks confirmed that they store their DHI under argon, in the dark, at 4°C. NB: Differences in DHI potency observed between cell-free and cellular assays are not unexpected, and are likely a consequence of several unavoidable factors. First, the reaction between DHI and Nurr1 is redox sensitive. The kinetics and thermodynamics of the interaction will likely differ between cell-free (non-reducing) and cellular (highly reducing) environments. Second, DHI/IQ is an unstable molecule that readily polymerizes with itself (and, in cells, with other molecules), reducing the effective concentration of the active ligand over time.

### Differential Scanning Fluorimetry Assays

The Nurr1 LBD protein was buffer exchanged from Base Buffer into 25 mM HEPES, 150 mM NaCl, pH 7.4 using a Zeba Spin Desalting Column (ThermoFisher, product number 89882). The DSF assays were carried out in a final volume of 50 μL, comprised of 4 μM protein, 2.5x SYPRO Orange (ThermoFisher/Life Technologies, from 5000x stock), the specified concentration of ligand in DMSO, and reaction buffer comprised of 25 mM HEPES, pH 7.4, 150 mM NaCl (2% final DMSO for each reaction). Samples were allowed to incubate in the dark for 30 min at 25 °C, prior to exposure to thermal gradient. Fluorescence was monitored using the ViiA 7 Real Time PCR System (ThermoFisher) in a 96 well polypropylene plate (Agilent Technologies Cat #410088) using a 2 MicroAmp Optical Adhesive Film, PCR compatible (P/N 4311971). The filter used for fluorescent monitoring corresponded to an excitation/emission of 470 ± 15 and 558 ± 11 nm, respectively. The thermal gradient was executed from 25 °C to 95 °C at a rate of 0.05 °C/s. Melting temperatures were determined based on the temperature corresponding to the maximum of derivative of the fluorescence signal (temperature vs dRFU/dT).

### Surface Plasmon Resonance Assays

Details for each experiment are summarized in the table below. Data were collected using either a Biacore T100 (GE) or Biacore T200 (GE) instrument at a flow rate of 30 μL/min and at a temperature of 25°C. The running buffer used in each experiment was either SPR Buffer A: 25mM HEPES, pH 7.4, 150mM NaCl, 0.05% Surfactant P20, and 2% DMSO, or SPR Buffer B: 10mM HEPES, pH 7.4, 150mM NaCl, 0.005% Surfactant P20, 3 mM EDTA, and 2% DMSO (diluted from 10x HBS-EP+; GE Healthcare Life Sciences). The biotinylated Nurr1 LBD was immobilized on a CAP chip (GE Healthcare Life Sciences; product number 28920234). Data collection was performed with one of the following three approaches: (i) no surface regeneration in between injections, (ii) surface regeneration between injections using 6 M guanidium HCl + 0.25 M NaOH regeneration solution followed by re-immobilization of Nurr1 (as described in the manufacturer’s regeneration protocol), or (iii) in single-cycle kinetics mode. DHI, dopamine, and L-DOPA dilutions were carried out starting from DMSO stocks (5 or 10 mM) in DMSO, whereas L-DOPA dilutions were carried out starting from a water stock (5 mM) in water. Analytes dissolved in DMSO were added to 1.02x running buffer without DMSO to yield a final DMSO concentration of 2%. Data processing included double referencing (i.e. reference flow cell and buffer subtracted using a buffer injection of appropriate contact time for the given injection), with solvent correction deemed unnecessary. For the single cycle kinetics approach, DHI was injected at four increasing concentrations (0.62, 1.25, 2.50, 5.00 μM) and responses were normalized to the initial Nurr1 LBD immobilized (summarized below). The surface was regenerated and fresh Nurr1 LBD was immobilized in between single-cycle kinetics experiments.

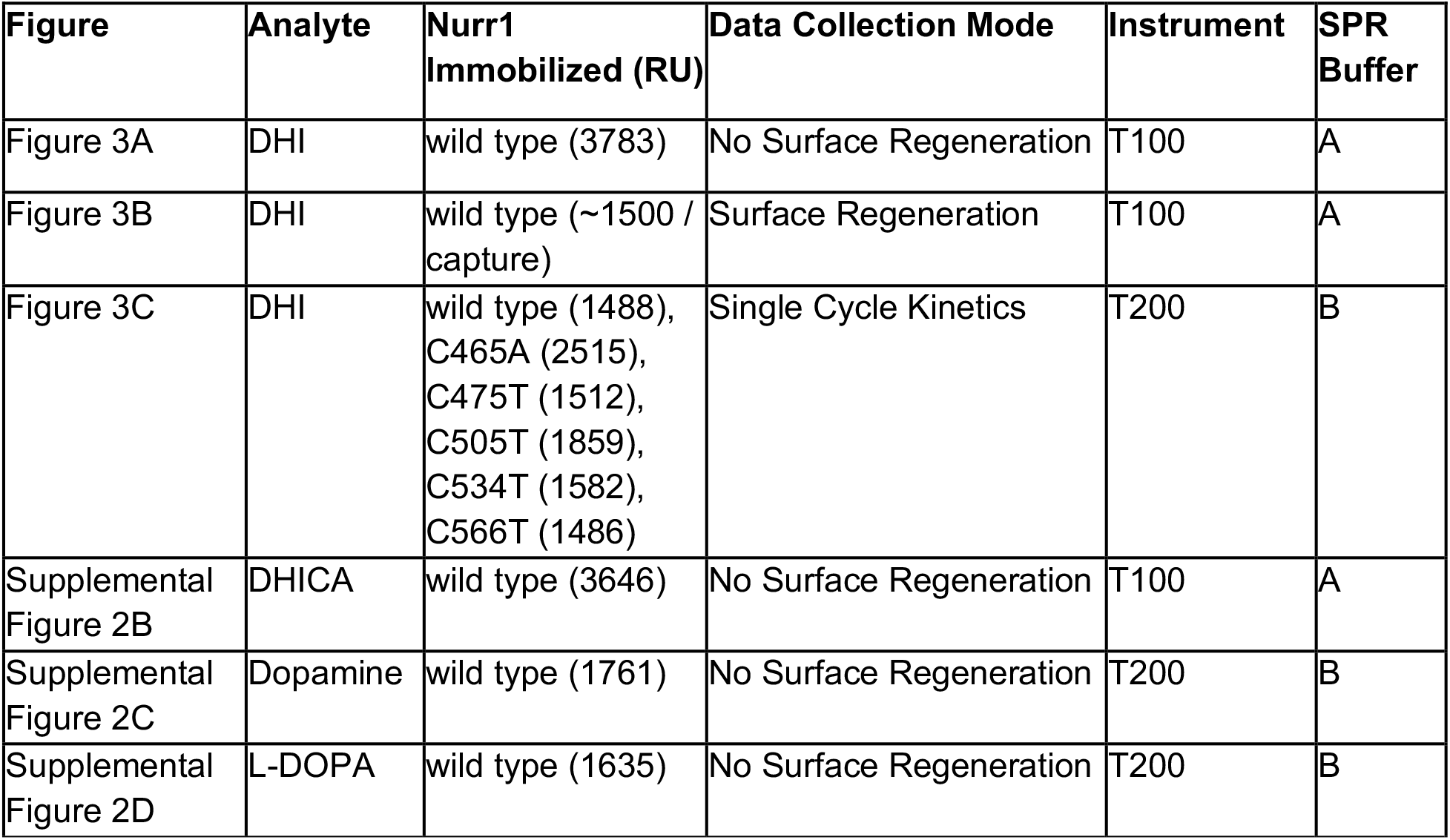

| Figure | Analyte | Nurr1 Immobilized (RU) | Data Collection Mode | Instrument | SPR Buffer |
| --- | --- | --- | --- | --- | --- |
| Figure 3A | DHI | wild type (3783) | No Surface Regeneration | T100 | A |
| Figure 3B | DHI | wild type (~1500 / capture) | Surface Regeneration | T100 | A |
| Figure 3C | DHI | wild type (1488),<br>C465A (2515),<br>C475T (1512),<br>C505T (1859),<br>C534T (1582),<br>C566T (1486) | Single Cycle Kinetics | T200 | B |
| Supplemental Figure 2B | DHICA | wild type (3646) | No Surface Regeneration | T100 | A |
| Supplemental Figure 2C | Dopamine | wild type (1761) | No Surface Regeneration | T200 | B |
| Supplemental Figure 2D | L-DOPA | wild type (1635) | No Surface Regeneration | T200 | B |

#### Determining Ligand Stoichiometry

The theoretical Rmax for DHI binding to the primary site (n = 1), and additionally to the secondary site (n = 2), in the Nurr1 LBD was calculated using the molecular weights of DHI (analyte) and the Nurr1 LBD (ligand), and the amount (RU) of immobilized Nurr1 captured to the chip (as described in the Biacore Handbook available online). The general equation is as follows:

Rmax (RU) = (analyte MW / ligand MW) x (immobilized ligand level in RU) x analyte stoichiometry

For our assays (see Figure 3B), the theoretical Rmax for one and two molecules of DHI bound per Nurr1 LBD was calculated as follows:

(n = 1 DHI/IQ bound) Rmax = (A / B) x (C x 1)
(n = 2 DHI/IQ bound) Rmax = (A / B) x (C x 2)
where A = 150 Da for DHI; B = 30632 Da and 30630 Da for the wildtype and cysteine point mutant Nurr1 LBDs, respectively; C = RU for Nurr1 immobilized in each experiment (see above).

### Crystallography of the Nurr1 LBD

#### Crystallization

The Nurr1 LBD was buffer exchanged into 150 mM NaCl, 25 mM HEPES, pH 7.4 using a 7 kD cutoff Zeba Desalting Column. DHI was then added to a final concentration of 200 μM (2% DMSO) from a 10 mM stock to a protein solution of 228 μM. The reaction was allowed to proceed on ice for 15 min before excess DHI and DMSO were removed via buffer exchange as described above. This labeling procedure was then repeated for a second time. Crystals were then prepared using a custom screen surrounding the previously reported crystallization condition (18% PEG3350, 200 mM KBr, 100 mM HEPES, pH 6.5)(Wang et al., 2003), with a pH range of 6.5-8.0 generated by mixing various volumes of 100 mM HEPES at pH 6.5 and 8.0. Crystals grew in a 96-well hanging drop format (1 drop per well) as 500 nL drops consisting of a 1:1 ratio of protein to condition with initial protein concentrations ranging from 4-7 mg/mL. Crystals were transferred briefly into the crystallization condition supplemented with 20% ethylene glycol to serve as a cryoprotectant prior to plunging into liquid nitrogen. The best diffracting crystal described herein was ~200 μm x 50 μm x 50 μm in size and cyroprotected 30 days after the initial drops were set.

#### Data Collection

Data was collected as the Advanced Light Source, Beamline 8.3.1 at an energy of 11111eV on a ADSC Quantum 315r S/N 926 detector under a stream of liquid nitrogen.

#### Structure Determination

Data were integrated and scaled using XDS (Kabsch, 2010). Although these crystals grew isomorphous to the deposited apo structure (PDB: 1OVL), crystals can reportedly grow as either the P3_1_ or P3_1_2_1_ space group (Wang et al., 2003). Exhaustive attempts at reaching an initial molecular replacement solution using the previously deposited apo Nurr1 LBD structure were unsuccessful, regardless of which space group was chosen for processing. We were initially able to refine this model using phenix.refine (Afonine et al., 2012) using the hexamer from 1OVL after applying the appropriate P3_1_ reindexing operator (-h,-k,l or - k,-h,-l). However, the Rwork-Rfree gap was ~10% after many rounds of refinement (and could not be closed), indicating that the higher symmetry P3_1_2_1_ might be the appropriate space group. We utilized Zanuda (Lebedev and Isupov, 2014) to help explore the higher symmetry P3_1_2_1_ as a potential space group with a trimer of Nurr1 monomers as the asymmetric unit. Zanuda successfully identified one potential trimer that could serve as an initial model for refinement. The data were then merged as P3_1_2_1_ using this trimer (chains A, B, and C of PDB: 1OVL) as a reference with the program *pointless* (Evans, 2011) followed by scaling with *aimless* (Evans and Murshudov, 2013). Iterative rounds of phenix.refine and model building with COOT (Emsley et al., 2010) were able to refine the model to within reasonable structure validation metrics. Chains A, B, and C of PDB: 1OVL were also used as a reference model for refinement. After several rounds of refinement and rebuilding, the Cys-IQ C2-adduct was introduced at Cys566 for chains A, B, and C using computational modeling as described below. TLS refinement and target-weight optimization for both XYZ and ADP restraints were performed. Occupancies were refined for the Cys-IQ C2-adducts (DHI/IQ atoms only, not Cys566 atoms) to allow for only partial adduct formation. Finally, phenix.polder was used to calculate difference maps for the adduct (Liebschner et al., 2017).

#### Initial Model Building of IQ Covalently Bound to Cys566

A partially refined structure without the Cys566-IQ adduct built in was used as a starting structure for computational modeling. The unnatural amino acid residue Cys566-IQ (N-methylated and C-acetylated) linked to IQ at the C2 atom was built within Maestro within the Schrodinger Suite. Hetgrp_ffgen (Schrodinger utilities) was then used to generate parameters for this adduct. The program PLOP (Jacobson et al., 2004) was then used to perform covalent docking for this cysteine adduct using 500,000 conformations followed by energy minimization of the introduced adduct. This model was then further refined with phenix.refine using a ligand restraint file for the Cys-IQ adduct (described below).

#### Ligand Restraint Generation

The Cys566-IQ unnatural amino acid was exported from Maestro in SMILES format and opened in JLigand. Atom names were manually modified to match the pdb atoms from the model generated using PLOP. This structure was saved as a .cif file and used as a template for phenix.elbow (Moriarty et al., 2009) ligand restraint generation using AM1 optimization.

### Positional Scanning Simulations

To explore the positional preference for thiolate attack, potential energy surfaces for the catechol and the quinone forms of DHI were scanned. All quantum mechanics calculations were carried out by using the Gaussian 09 software (Frisch et al., 2016). Geometries were optimized at the B3LYP/6-311+G(d) level.

### Luciferase Assays

Human choriocarcinoma JEG-3 cells were maintained in Eagle’s MEM with Earle’s BSS (MEM) supplemented with 10% FBS, 1% NEAA, 1% L-Glutamine, 1% Sodium Pyruvate and 1% Penicillin/Streptomycin. For transfection, cells were seeded in 96-well plate in antibiotic free medium one day before transfection. Transfections were carried out with Lipofectamine 2000 (Invitrogen) according to manufacturer’s protocol. Lipofectamine/DNA complexes were prepared in Opti-Mem medium and incubated with cells overnight. The Nurr1 LBD fragment was subcloned into pM plasmid (Clontech) containing GAL4 DNA binding domain. The reporter plasmid, pGL4.35 (luc2P/9XGAL4 UAS/Hygro) Vector (Promega), contains 9 repeats of GAL4 UAS (Upstream Activator Sequence), which drives transcription of the luciferase reporter gene luc2P in response to binding of Gal4 DNA binding domain fused protein. The pRL-TK (Promega) plasmid expressing renilla luciferase was used as an internal control. The amounts of pM-Nurr1-LBD, pGL4.35 and pRL-TK were 50 ng, 50 ng and 5 ng per well respectively. Six hours before luciferase measurement, different doses of DHI were added and incubated. Cells from each well were lysed with 100 μl Passive Lysis Buffer (Promega). The luciferase activities of firefly and renilla were measured using Dual-Luciferase^®^ Reporter Assay (Promega) according to manufacturer’s protocol. The results were expressed as average of firefly/renilla luciferase activity ± SEM (n = 4).

### Target Gene Transcription Assays

#### DHI Administration

Larval zebrafish (Danio rerio) used for experiments were bred in house and treated in accordance with IACUC regulations. Fifty zebrafish larvae per concentration per experiment, for a total of 200 larvae per concentration (replicates = 4) were treated in a 10 cm Petri dishes at 72 h post fertilization (hpf) with DHI diluted (100 mM stock) into 40 mL of blue egg water (lab stock) to reach final concentrations of 10 and 100 μM (0.2% DMSO). Control larvae were exposed to an equivalent amount of DMSO (0.2%). Samples were collected for further analysis 6 and 24 h post treatment.

#### RNA Isolation and Transcript Quantification

Total RNA was extracted from zebrafish larvae (collection post-treatment at 78 hpf after 6 h DHI exposure and 96 hpf after 24 h DHI exposure) using TRIzol reagent (Invitrogen) by homogenization and purified using RNeasy Mini Kit (Qiagen). The cDNA was then synthesized from 160 ng of purified RNA using qScript cDNA SuperMix (Quanta Biosciences) and used as template. The qPCR was performed using Applied Biosystems SYBR Green PCR Master Mix and the ABI7900HT machine. The forward and reverse primers (Supplemental Table 5) were designed using NCBI/primer-BLAST software with exon-exon junction parameters and *Danio rerio* RefSeq for off targets. The elongation factor 1 alpha (*elf1α)* gene was used as internal housekeeping gene to determine the relative mRNA expression. The *elf1α* primers were used as standard control (McCurley and Callard, 2008) to generate Ct values. Transcript levels for target genes were normalized to the housekeeping gene *elf1a* and fold change was compared to gene expression levels from DMSO-treated larvae. GraphPad Prism 7 software was used for statistical analysis. Unpaired Student t-test was applied for DMSO ct *vs* 10 μM and 100 μM DHI. Selected statistical significance is defined as follows: P > 0.05 ns, P ≤ 0.05*; P ≤ 0.01**; P ≤ 0.001***; P ≤ 0.0001****. Significant results (P values) were observed for the following conditions: 6 h 100 μM DHI treatment (*vmat2* = 0.019; *dat* = 0.015) and 24 h 100 μM DHI treatment (*th =* 0.030 and *vmat2* = 0.00008).

## Author Contributions

Biophysical assays, molecular modeling, and crystallography were performed by J. B.; quantum mechanical calculations were performed by B. T.; cell-based assays were performed by Y. W. and P. B.; preparation of mutant protein was performed by S. K.; zebrafish assays were performed by F. O. and H. L.

## Acknowledgements

This work was supported by grants from the Roger’s Family Foundation, the CTSI, the MAKK Seed Fund, the NIH (R01 NS108404-01), and the Program for Breakthrough Biomedical Research that is partially funded by the Sandler Foundation. We thank Joseph Wagner for helpful discussions. Beamline 8.3.1 at the Advanced Light Source is operated by the University of California Office of the President, Multicampus Research Programs and Initiatives grant MR-15-328599 the National Institutes of Health (R01 GM124149 and P30 GM124169), Plexxikon Inc. and the Integrated Diffraction Analysis Technologies program of the US Department of Energy Office of Biological and Environmental Research. The Advanced Light Source (Berkeley, CA) is a national user facility operated by Lawrence Berkeley National Laboratory on behalf of the US Department of Energy under contract number DE-AC02-05CH11231, Office of Basic Energy Sciences.

---

**Supplemental Figure 1.**
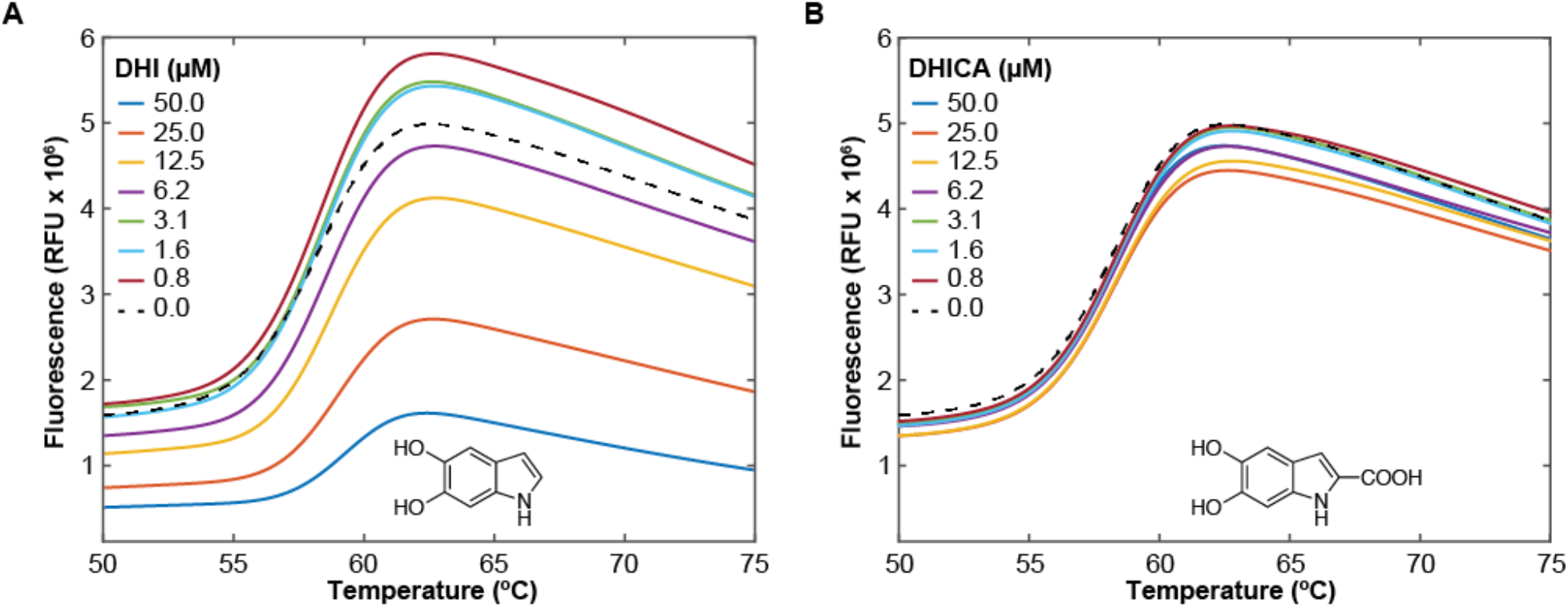
DHI but not DHICA decreases the overall SYPRO orange fluorescence. DSF assays were carried out on the Nurr1 LBD (4 μM) in the presence of the dye SYPRO orange (2.5x) and varying concentrations of DHI or DHICA. **(A, B)** Plot of the fluorescence (RFU) as a function of temperature for Nurr1 in the presence of increasing concentrations of (A) DHI and (B) DHICA. Experiments were performed in triplicate.

**Supplemental Figure 2.**
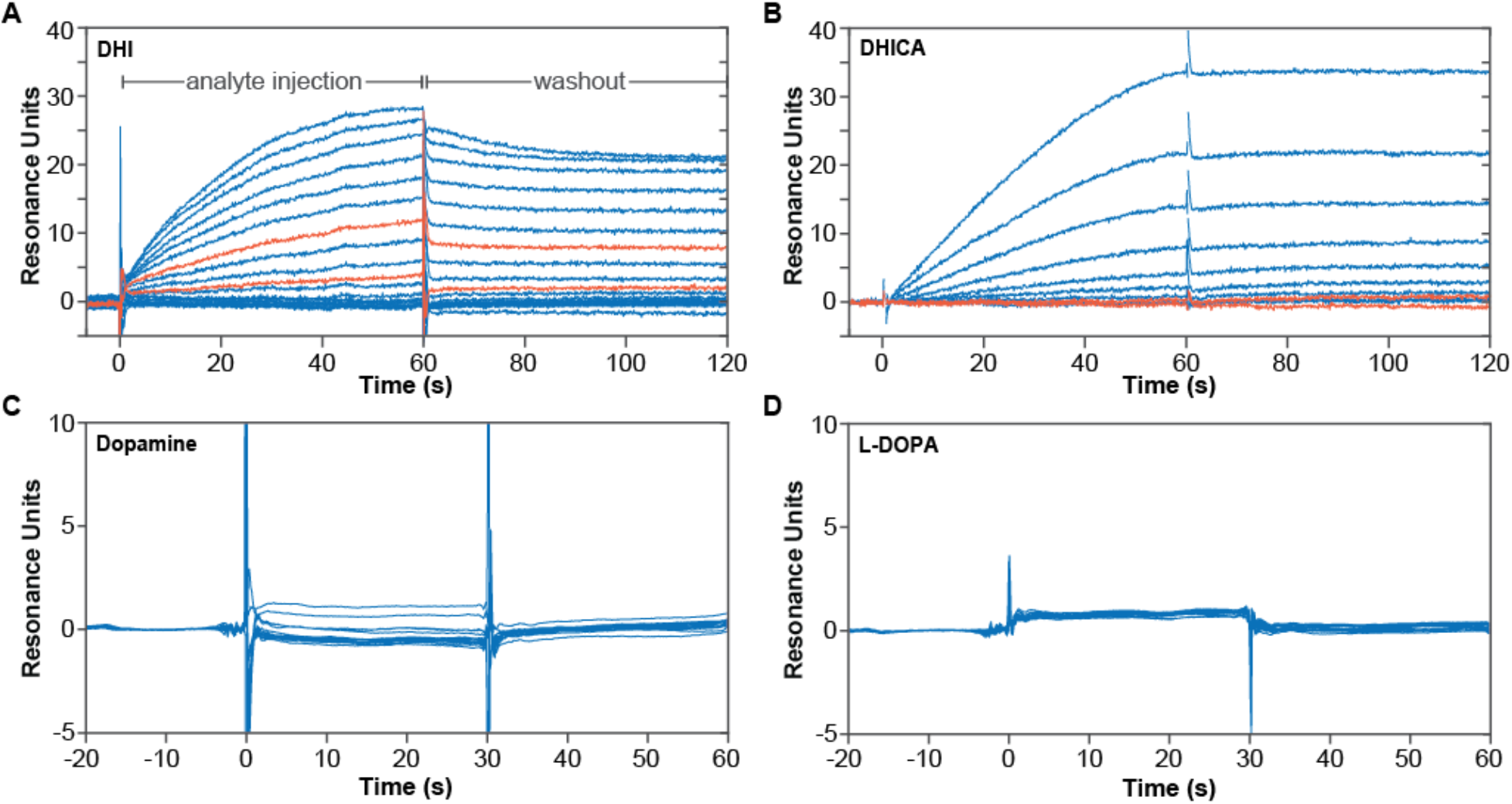
Specific association of DHI, but not DHICA, dopamine or L-DOPA, with the Nurr1 LBD measured by surface plasmon resonance. Biotinylated Nurr1 LBD was coupled to a streptaviden-coated biosensor chip and exposed to analytes at varying concentrations without surface regeneration between exposures. **(A)** Concentration-dependent association of DHI with Nurr1 LBD. DHI was injected at 15 increasing concentrations (16.8 μM DHI, 0.6x dilutions down to 0.01 μM; 60 s association time). Sensorgrams for DHI injections at 0.17 and 0.78 μM are highlighted in orange. **(B)** Non-specific association of DHICA with Nurr1 LBD. DHICA was injected at 11 increasing concentrations (25.9 μM DHICA, 0.6x dilutions down to 0.16 μM; 60 s association time). Sensorgrams for DHICA injections at 0.16 and 0.73 μM are highlighted in orange. **(C, D)** No association of dopamine or L-DOPA with Nurr1 LBD. Analytes were injected at 16 increasing concentrations (100 μM dopamine or L-DOPA, 0.6x dilutions down to 0.05 μM; 30 s association time).

**Supplemental Figure 3.**
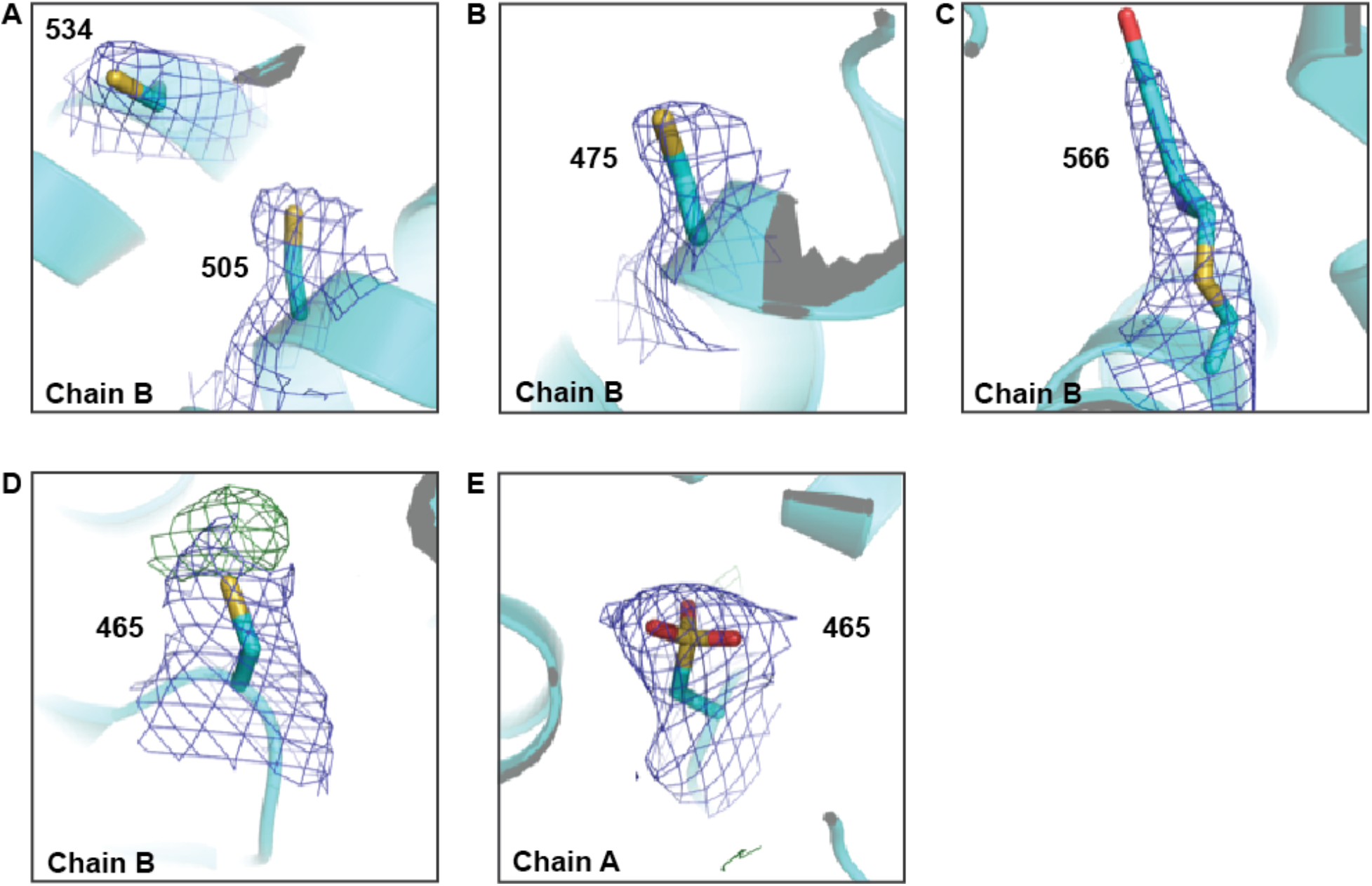
Electron density of all cysteine residues in the Nurr1 LBD-IQ structure. Electron density map (2mFo-DFc; contoured to 1σ) is shown in blue mesh. Difference density map (mFo-DFc; contoured to 3σ) is shown in green mesh. Cysteines 534, 505, and 475 show no positive density around their respective sulfur atoms in all three chains (data for Chain B shown), whereas Cys566 shows positive electron density around its sulfur atom that is consistent with the IQ-adduct in each chain. Cysteine 465 shows positive electron density around the sulfur atom in Chain B and Chain A. However, an IQ adduct at this position is precluded by severe steric clashes. Rather, the electron density is satisfied by modeling oxidized cysteine (sulfonic acid) at this position in Chain A (70% occupancy), but not Chain B, using occupancy refinement; accordingly, Cys465 of Chain B still shows positive electron density around its sulfur atom. Notably, Cys465 is near two arginine residues (Arg450 and Arg454) that could favorably interact with oxidized cysteine.

**Supplemental Figure 4.**
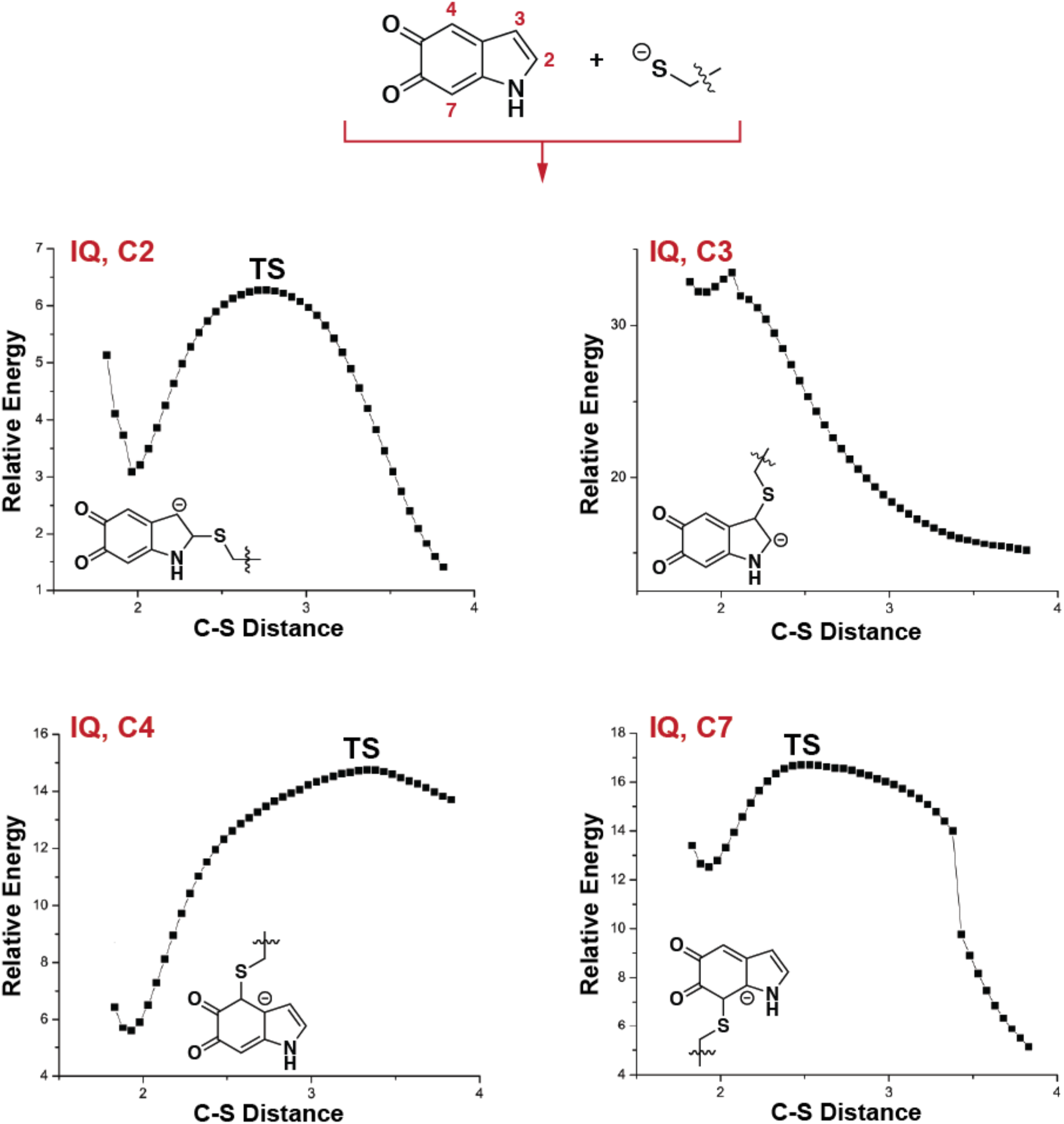
Potential energy surface for thiolate attack on IQ (indolequinone) at each of four potentially reactive sites (C2, C3, C4, C7). The reactions at positions C2, C4, and C7 exhibit an observable energy barrier, 6.3, 14.7 and 16.7 kcal/mol respectively, suggesting the existence of a stable C-S linked reaction product. The low reaction energy barrier observed for the reaction at C2 suggests reversibility. C-S distances are in Å; relative energies are in kcal/mol; TS = transition state.

**Supplemental Figure 5.**
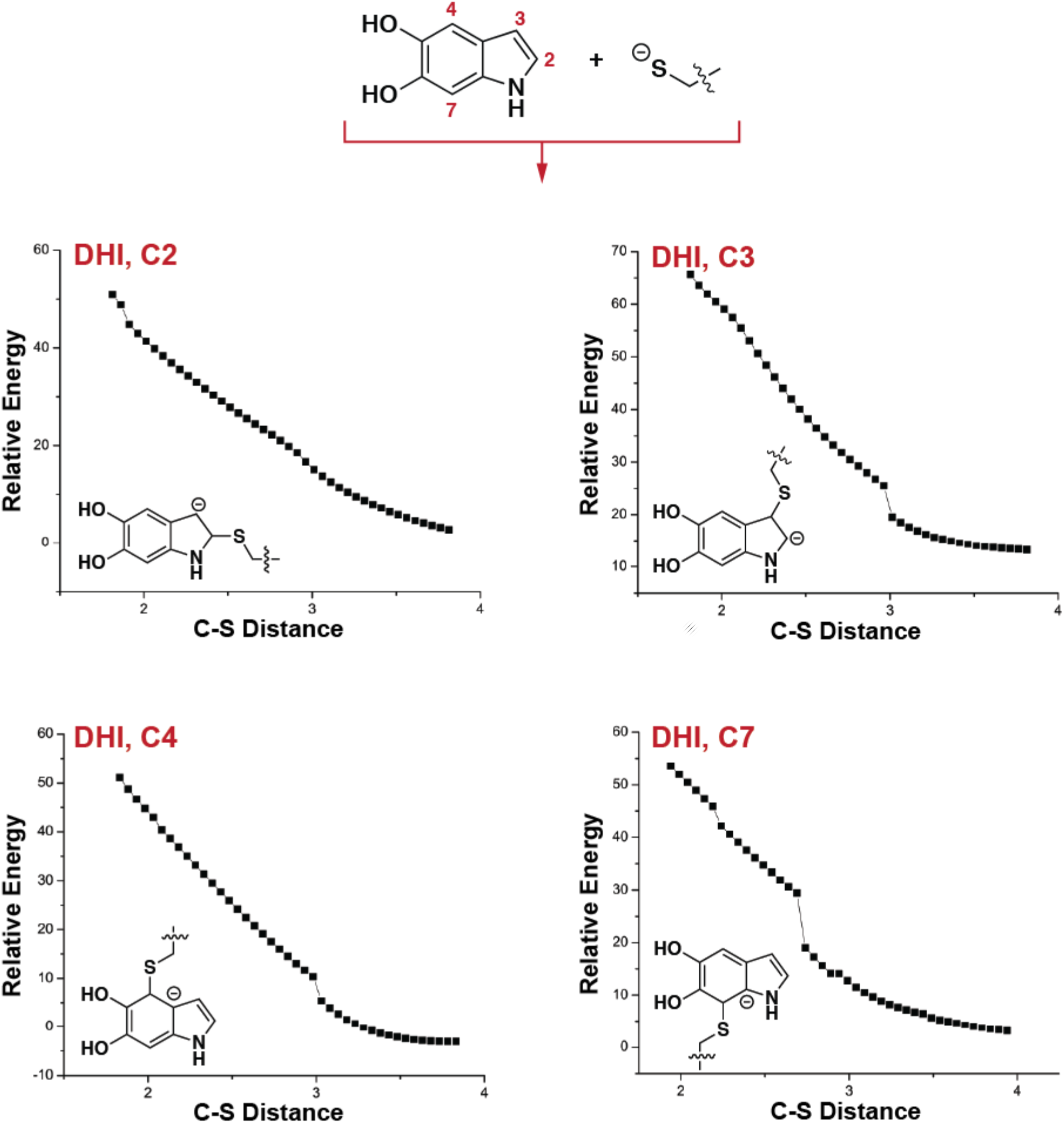
Potential energy surface for thiolate attack on DHI (catechol) at each of four potentially reactive sites (C2, C3, C4, C7). None of these reactions exhibit an energy barrier that would indicate formation of a stable C-S bond at any site. C-S distances are in Å; relative energies are in kcal/mol.

**Supplemental Figure 6.**
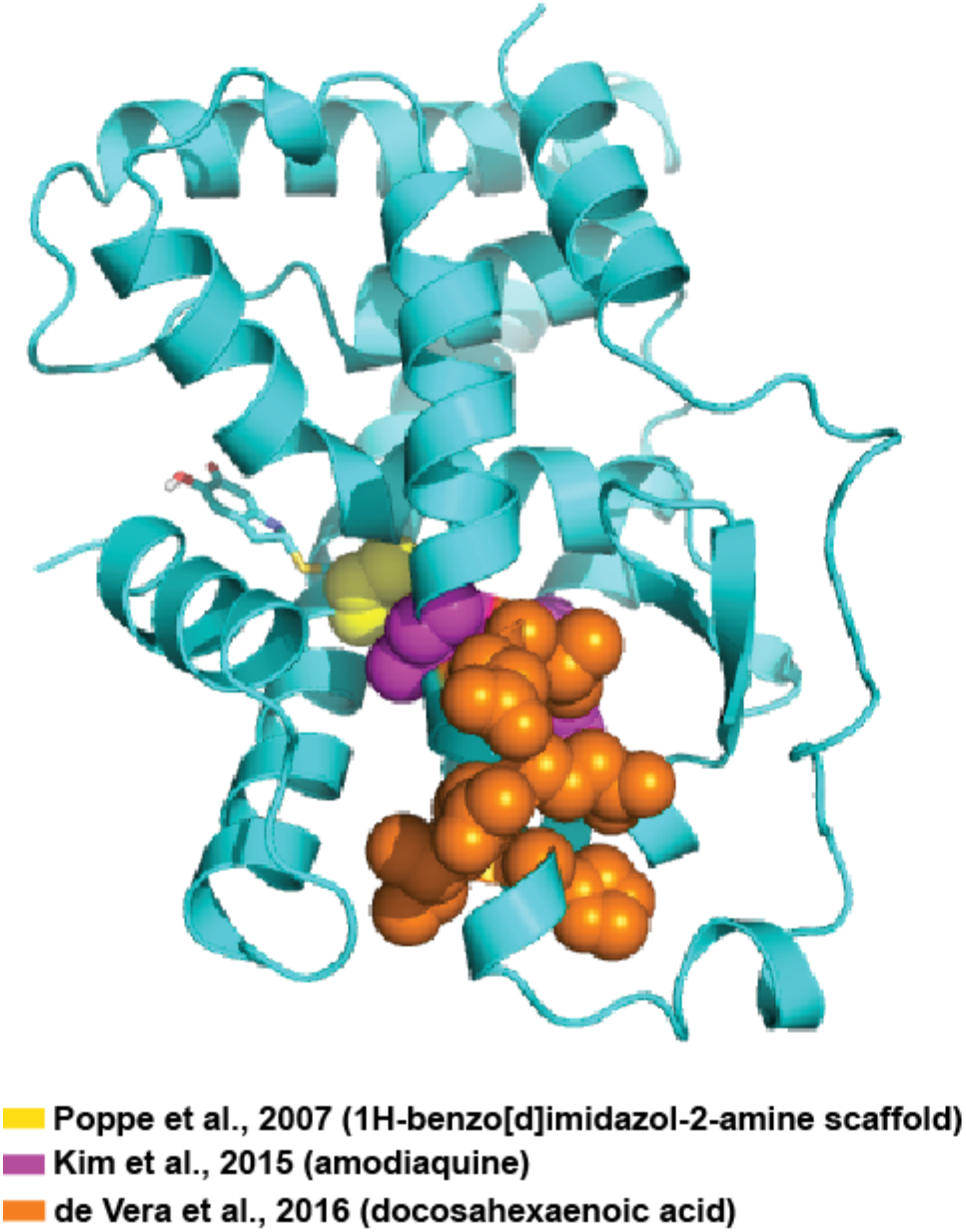
The DHI/IQ binding site is distinct from the amodiaquine and docosahexaenoic acid binding sites. IQ adduct shown as sticks. Amino acid residues implicated in binding other ligands by NMR spectroscopy are shown as spheres. Poppe et al. identified Cys566 (yellow spheres) as part of the binding site for benzimidazole-like compounds. Kim et al., identified residues (magenta spheres) forming a non-canonical binding site for amodiaquine. de Vera et al., identified residues (orange spheres) forming a canonical binding site for fatty acids.

**Supplemental Table 1.**
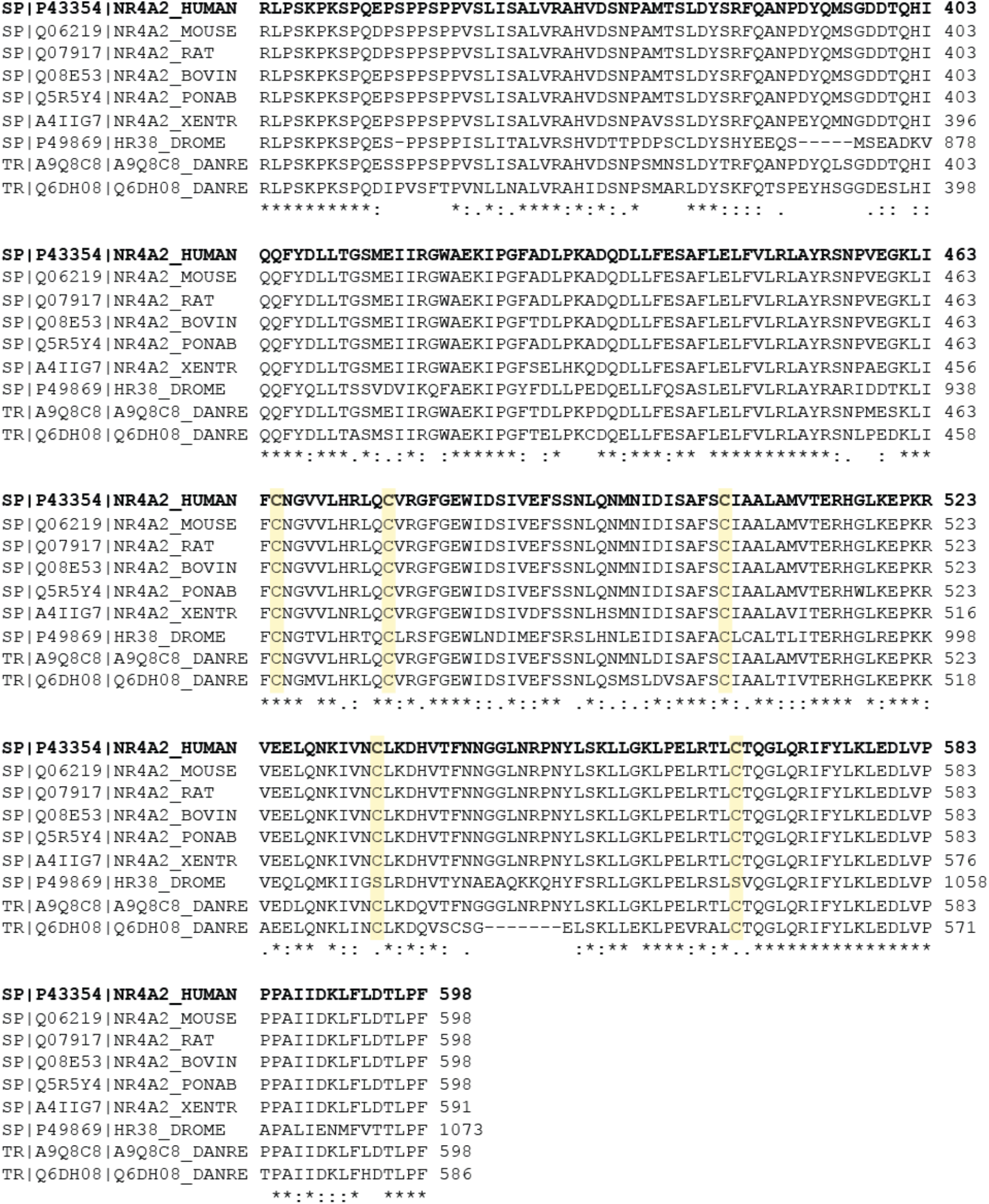
Five cysteine residues within the Nurr1 LBD are conserved across Nurr1 orthologs. Multiple sequence alignment was performed using the Clustal Omega server. Cysteine residues are highlighted in yellow.

**Supplemental Table 2.** Data collection and refinement statistics.

|  | **Nurr1-DHI/IQ<sup>#</sup>** |
| --- | --- |
| **Wavelength** | 1.116 |
| **Resolution range** | 40.29 - 3.2 (3.314 - 3.2) |
| **Space group** | P 31 2 1 |
| **Unit cell** | 80.586 80.586 225.805 90 90 120 |
| **Total reflections** | 169966 (16629) |
| **Unique reflections** | 14697 (1414) |
| **Multiplicity** | 11.6 (11.7) |
| **Completeness (%)** | 99.58 (99.58) |
| **Mean I/sigma(I)** | 13.74 (1.22) |
| **Wilson B-factor** | 98.95 |
| **R-merge** | 0.201 (2.643) |
| **R-meas** | 0.2103 (2.762) |
| **R-pim** | 0.06123 (0.7986) |
| **CC1/2** | 0.999 (0.561) |
| **CC\*** | 1 (0.848) |
| **Reflections used in refinement** | 14647 (1413) |
| **Reflections used for R-free** | 1445 (140) |
| **R-work** | 0.2489 (0.4005) |
| **R-free** | 0.2954 (0.4024) |
| **CC(work)** | 0.962 (0.614) |
| **CC(free)** | 0.901 (0.527) |
| **Number of non-hydrogen atoms** | 5529 |
| **macromolecules** | 5517 |
| **ligands** | 12 |
| **Protein residues** | 685 |
| **RMS(bonds)** | 0.011 |
| **RMS(angles)** | 1.47 |
| **Ramachandran favored (%)** | 96.05 |
| **Ramachandran allowed (%)** | 3.49 |
| **Ramachandran outliers (%)** | 0.46 |
| **Rotamer outliers (%)** | 0.99 |
| **Clashscore** | 24.39 |
| **Average B-factor** | 120.83 |
| **macromolecules** | 120.85 |
| **ligands** | 111.03 |
| **Number of TLS groups** | 13 |

<sup>#</sup>Statistics for the highest-resolution shell are shown in parentheses.

**Supplemental Table 3.** Polder map statistics for Cys566-IQ adduct. Only IQ atoms (i.e. no Cys566 atoms) were used in the calculations.

|  |  |
| --- | --- |
| CC(1,2) | 0.6288 |
| CC(1,3) | 0.7835 |
| CC(2,3) | 0.6191 |
| Peak CC(1,2) | 0.6465 |
| Peak CC(1,3) | 0.7748 |
| Peak CC(2,3) | 0.6200 |

**Supplemental Table 4.**
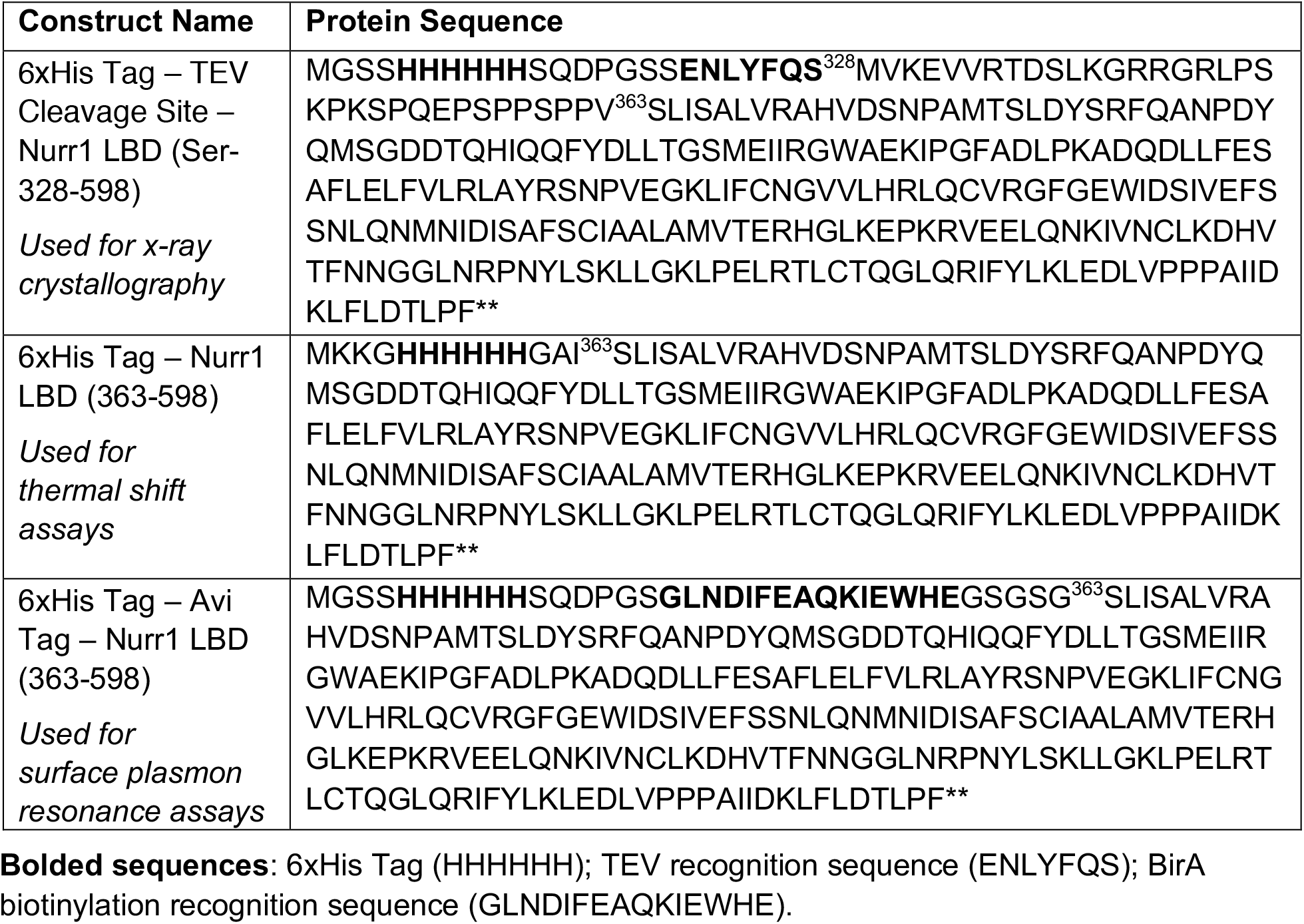
Nurr1 LBD construct names, corresponding protein sequence, and associated experiments.

**Supplemental Table 5.**
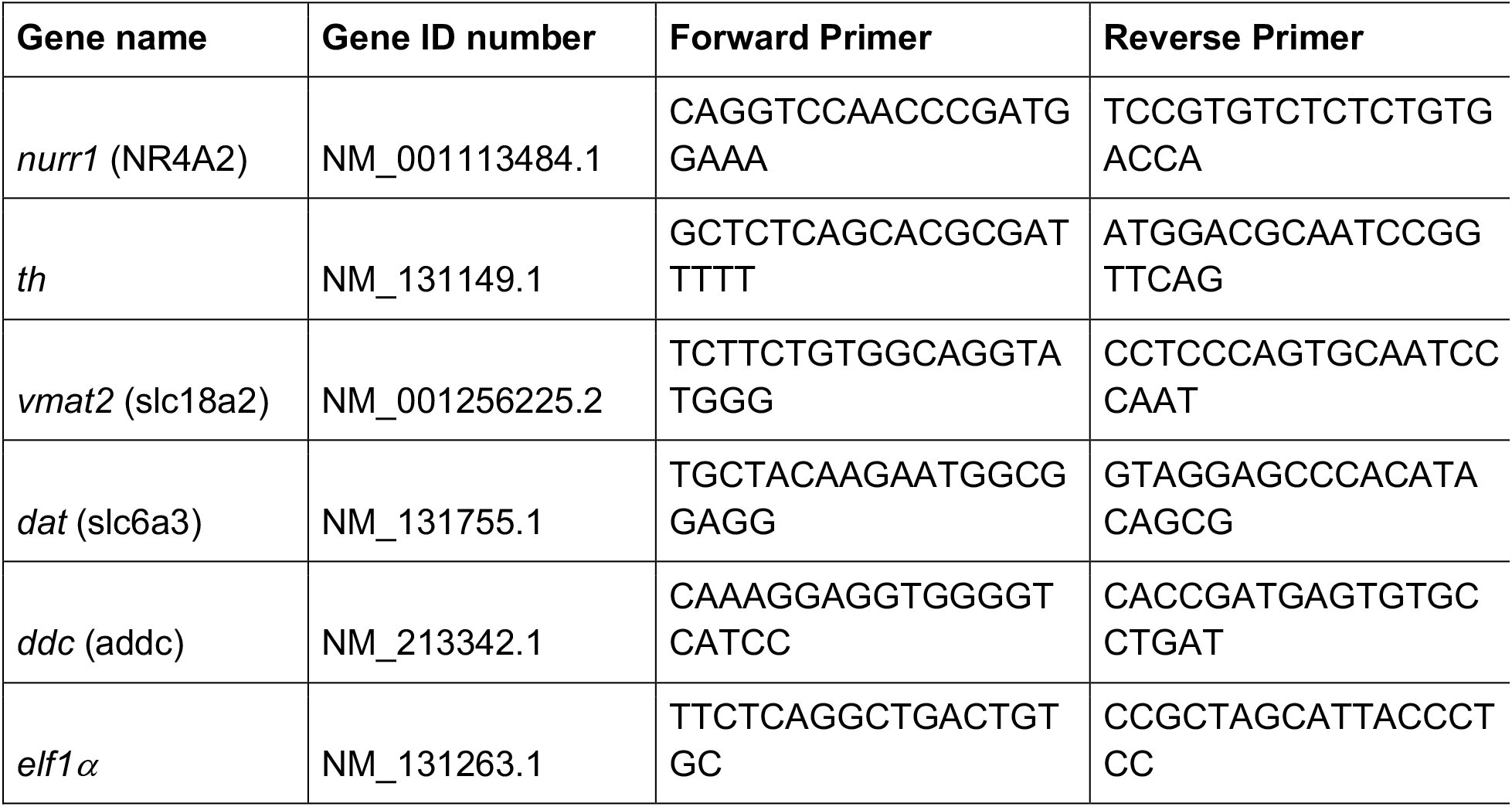
Primers used for qPCR.

## Footnotes

1 Rmax is the maximum analyte (DHI) binding capacity of the immobilized ligand (Nurr1 LBD) expressed in resonance units (RU); see Experimental Procedures for additional information.

2 The term “canonical ligand binding pocket” refers to the expected site of ligand binding based on the classic binding pocket for ligands in well-characterized NRs, such as androgen, estrogen, and glucocorticoid receptors.

3 A cation-pi interaction between arginine and tryptophan (indole) residues is frequently observed in structural biology; over one-fourth of the tryptophans in the protein data bank experience an energetically significant cation-pi interaction (see Gallivan and Dougherty, 1999).

4 We assume that the promoter sequences in zebrafish are similar to those identified in mammals.

5 In these examples, the specific modifications could not be unambiguously identified, as the reactive metabolites were generated by treating dopamine with tyrosinase or chemical oxidants (i.e. NaIO_4_, H_2_O_2_), resulting in the production of several different reactive species (e.g., see Figure 1B).

